# mTORC2 coordinates the leading and trailing edge cytoskeletal programs during neutrophil migration

**DOI:** 10.1101/2022.03.25.484773

**Authors:** Suvrajit Saha, Jason P. Town, Orion D. Weiner

## Abstract

By acting both upstream and downstream of biochemical organizers of the cytoskeleton, physical forces function as central integrators of cell shape and movement. Here we use a combination of genetic, pharmacological, and optogenetic perturbations to probe the role of the conserved mechanoresponsive mTORC2 program in neutrophil polarity and motility. We find that the tension-based inhibition of leading edge signals (Rac, F-actin) that underlies protrusion competition is gated by the kinase-independent role of the complex, whereas the mTORC2 kinase arm is essential for regulation of Rho activity and Myosin II-based contraction at the trailing edge. Cells required mTORC2 for spatial and temporal coordination between the front and back polarity programs and persistent migration under confinement. mTORC2 is in a mechanosensory cascade, but membrane stretch did not suffice to stimulate mTORC2 unless the co-input PIP_3_ was also present. Our work suggests that different signalling arms of mTORC2 regulate spatially and molecularly divergent cytoskeletal programs allowing efficient coordination of neutrophil shape and movement.

## INTRODUCTION

Directed cell migration underlies a wide range of physiological processes ranging from developmental morphogenesis to immune cell responses (SenGupta et al., 2021). Single cells move by extending a leading front that protrudes and a trailing rear that contracts and follows the front. These programs exhibit not only spatial compartmentalization of distinct intracellular signals to either the front or back of the cell (polarization) but also temporal coordination between these domains (Ku et al., 2012; Tsai et al., 2019; Xu et al., 2003). Neutrophils are a type of innate immune cell that rely on properly oriented cell polarity to migrate to sites of injury where they hunt and kill invading pathogens (Lämmermann et al., 2013; Liew and Kubes, 2019). In the cell front, activation of small GTPase Rac helps sets the permissive zone for WAVE-regulatory complex (WRC)-dependent actin polymerisation in protrusions (Koronakis et al., 2011; Lebensohn and Kirschner, 2009; Rottner et al., 2021; Srinivasan et al., 2003; Wang et al., 2002; Weiner et al., 2006). At the back, the GTPase RhoA stimulates myosin based contractility (Hind et al., 2016; Tsai et al., 2019; Wong et al., 2006). These signaling domains are sustained by short-range positive feedback loops within the modules and are spatially separated by mutual antagonism between them (Ku et al., 2012; Wang et al., 2013; Xu et al., 2003). Coordination within and between the modules is critical for polarity maintenance during persistent migration (Maiuri et al., 2015; Tsai et al., 2019; Yang et al., 2015), but how this coordination is achieved is not fully understood.

When cells protrude or contract, they alter the mechanical properties of the cell surface. While mechanics was initially seen as a downstream output of cytoskeletal dynamics, emerging evidence suggest that mechanics feeds back to regulate the upstream leading and trailing edge biochemical signals (Diz-Muñoz et al., 2013; Graziano et al., 2019; Hetmanski et al., 2019; Lieber et al., 2013; Mueller et al., 2017; Saha et al., 2018). In neutrophils, membrane tension acts as a long-range inhibitor of actin nucleation and polymerization to constrain the size and number of cell protrusions (Houk et al., 2012). Increases in membrane tension trigger a mechanosensitive signalling cascade to regulate actin dynamics in neutrophils. Actin-based polymerization in protrusions stimulates Mechanistic target of Rapamycin Complex 2 (mTORC2) through the activation of Phospholipase D2. By connecting increases in protrusion to decreases in actin nucleation, mTORC2 is a central component of the negative-feedback-based homeostat on membrane tension (Diz-Muñoz et al., 2016).

mTOR kinase is an ancient and evolutionarily conserved regulator of cell growth, proliferation, and survival (Saxton and Sabatini, 2017). mTOR forms two distinct multi-subunit complexes in mammalian cells mTOR complex 1 (mTORC1) and mTOR complex 2 (mTORC2). mTORC2 is formed from the association of core mTOR subunits with Rictor and mSin1. Rictor scaffolds the complex and is indispensable for the stability of the complex, whereas mSin1 aids the kinase activity of the complex (Frias et al., 2006; Jacinto et al., 2004; Sarbassov et al., 2004). mTORC2 activity is thought to broadly localize to plasma membrane (Berchtold et al., 2012; Ebner et al., 2017b; Riggi et al., 2020), where it relays growth factor signals by phosphorylating its downstream effectors Akt, PKC and SGK1 and other targets to regulate a wide range of cellular processes including cytoskeletal organisation and cell migration (Liu and Parent, 2011; Oh and Jacinto, 2011).

mTORC2 plays a homeostatic role in response to membrane stretch in a wide variety of cell types, ranging from yeast (Berchtold et al., 2012; Riggi et al., 2020, 2018) to immune cells (Diz-Muñoz et al., 2016) to *Dictyostelium* (Artemenko et al., 2016; Kamimura et al., 2008). In *S. cerevisiae*, plasma membrane (PM) stretch activates TORC2 to stimulate sphingolipid biosynthesis, sterol recycling and bilayer asymmetry as homeostatic mechanisms to reset membrane tension and restore membrane trafficking (Riggi et al., 2020; Roelants et al., 2017). In neutrophils, where membrane tension increases arise from protrusive forces of F-actin, mTORC2 based inhibition of actin polymerisation serves a mechanism to maintain the tension setpoint as well as restrict polarity signals (Diz-Muñoz et al., 2016; Houk et al., 2012; Liu et al., 2010). A conserved role of mTORC2 also involves gating chemoattractant signaling to cyclic- AMP production to regulate myosin contractility and drive tail retraction in neutrophils (Liu et al., 2014, 2010). The *Dictyostelium* homolog of Rictor, Pianissimo was initially identified in a genetic screen for regulators of chemotaxis before it was known to be part of TORC2 (Chen et al., 1997). Perturbation of mTORC2 component Rictor led to impaired chemotaxis in neutrophils, fibroblasts and *Dictyostelium* consistent with reports of its role in regulating actin cytoskeleton (Agarwal et al., 2013; He et al., 2013; Lee et al., 2005; Liu et al., 2010). These wide arrays of cytoskeletal defects are thought to rise from both positive (He et al., 2013) and negative inputs (Diz-Muñoz et al., 2016; Huang et al., 2017) to front and back polarity programs and have been hard to decouple.

Here, we investigate the relative contributions of the kinase dependent versus independent arms of Rictor/mTORC2 in regulating front and back polarity activation and coordination (**Fig 1A**). The kinase-independent arm of mTORC2 restricts actin polymerization at the cell front, whereas the kinase arm regulates myosin contractility at the cell back. Whereas front/back regulation are normally highly coordinated, they lose their coordination in the absence of mTORC2. These defects are particularly profound when neutrophils explore and move in confined environments. Stretch alone is not sufficient to activate mTORC2 unless the co-input of PIP_3_ is also present. Our work reveals a role for mTORC2 in coordinating front and back regulation through different effector arms of this highly-conserved mechanosensor.

**Figure 1.**
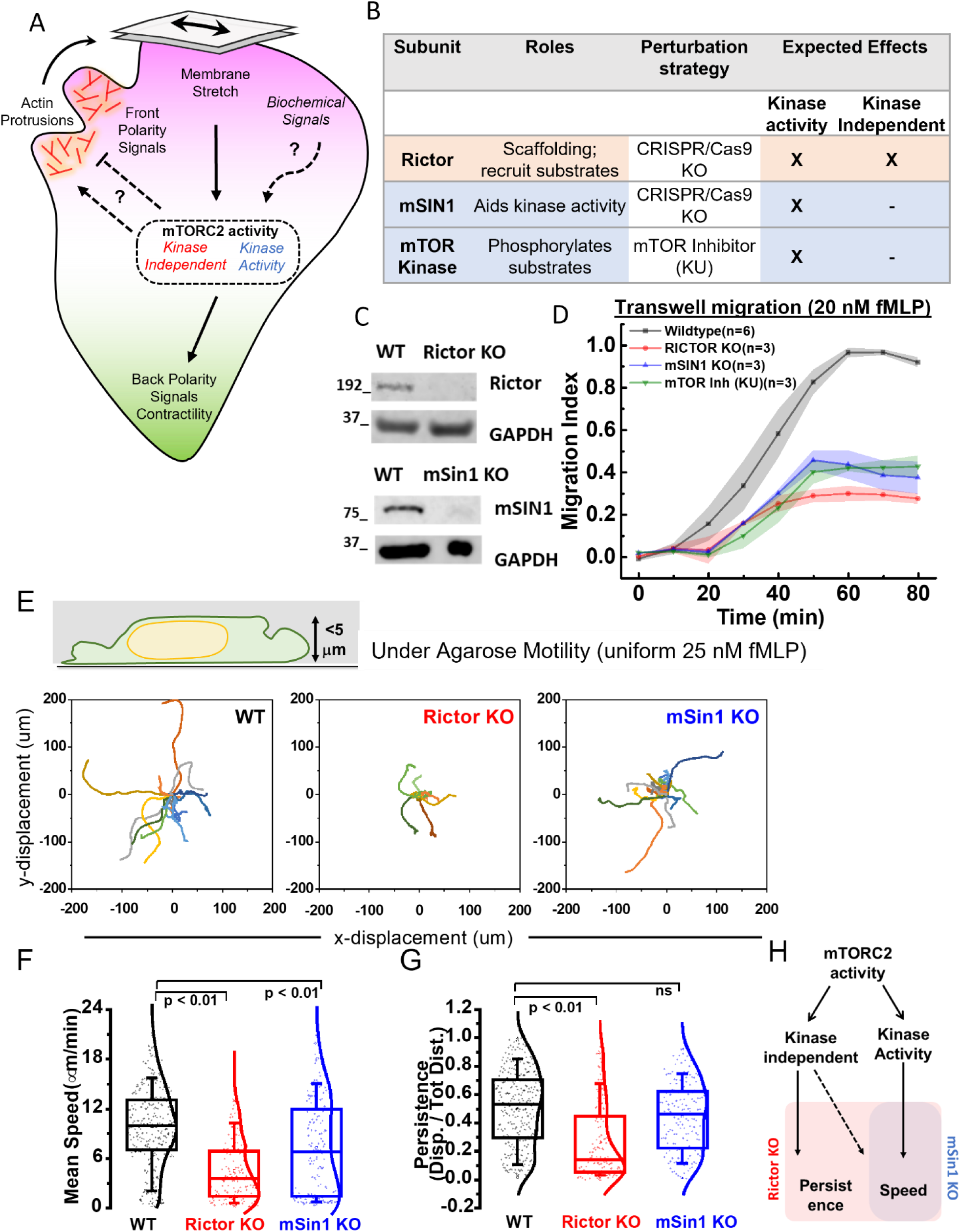
Rictor/mTORC2 is a mechanoresponsive regulator of neutrophil motility. (A) mTORC2 connects membrane stretch to regulation of front (magenta) and back (green) polarity programs, but how mTORC2 is activated (purely mechanical or requires biochemical co-inputs) and what aspect of mTORC2 activation (kinase-dependent vs independent roles) regulates these polarity signals is not understood. (B) To dissect the roles of kinase-dependent and kinase-independent roles of mTORC2, we generated individual CRISPR-Cas9 knockout lines of key components of the complex: Rictor (which scaffolds and aids structural integrity of the complex) or mSin1 (which primarily aids kinase activity). Additionally, mTOR Kinase inhibitors (here KU) would phenocopy mSin1 KO defects. (C) Representative immunoblots of wildtype (WT) HL-60 cells and gene-edited Rictor KO (top) and mSin1 KO (bottom) clonal HL-60 line to validate the loss of Rictor or mSin1 protein expression. GAPDH was used as a loading control. (D) Perturbation of mTORC2 activities in Rictor KO (n=3; red), mSin1KO(n=3; blue) and via mTOR Kinase inhibitor (KU; n=3; green) all led to defective transwell migration towards chemoattractant 20nM fMLP in comparison to WT cells (n=6; black). Mean ± SEM is plotted, n indicates independent replicates. (E) Schematic shows neutrophil-like dHL60 cell moving under an agarose (2%) overlay with uniform chemoattractant (25 nM fMLP). Randomly-chosen representative tracks (15 each) of wildtype (WT), Rictor KO, or mSin1KO cells over a 12 min observation window; axes show x-y displacement in μm. Rictor KO cells migrate poorly and have markedly shorter displacements. (F, G) Box plots (with kernel smooth distribution curve) show mean speed (F) and persistence (G; ratio of displacement/distance) averaged over individual tracks. Both Rictor KO and mSin1 KO cells shows a significant reduction (p < 0.01; one-way ANOVA with Tukey-means comparison) in migration speed compared to Wildtype. However, only Rictor KO show a significant decrease in the persistence (p < 0.01; one-way ANOVA with Tukey-means comparison). N = 294 (WT), 138 (RictorKO) and 165 (mSin1KO) tracks from individual cells pooled across 3 independent experiments. For box plots, median is indicated by the line, inter-quartile range (IQR) sets the box width and error bars indicate 10-90^th^ percentile. (H) Schematic highlights the phenotypes observed for mSin1KO and Rictor KO cells. Kinase-dependent roles of mTORC2 appear to regulate speed whereas kinase-independent role regulates both persistence and speed.

## RESULTS

### Rictor/mTORC2 is a mechanoresponsive regulator of neutrophil ameboid motility

How is mechanoresponsive triggering of TORC2 linked to its regulation of cell migrationOne possible route to regulating motility is through altering the dynamics of actin polymerisation or myosin contractility (**Fig 1A**). Earlier studies in neutrophils using partial knockdown of Rictor or mSin1 (using shRNAs) have found both positive and negative roles of TORC2 on cytoskeletal effectors (Rac, Cdc42, RhoA) confounding a clear understanding of the logic of this regulation (Diz-Muñoz et al., 2016; He et al., 2013; Liu et al., 2010). We reasoned genetic nulls with complete loss of mTORC2 specific subunits in neutrophil-like dHL60 cells would offer a more surgical approach to dissect these mechanisms, following the success of this strategy in other cell-based models (Agarwal et al., 2013; Guertin et al., 2006; Huang et al., 2017).

To distinguish between the relative contribution of the kinase roles of mTORC2 and the kinase-independent scaffolding roles of Rictor, we devised a CRISPR-Cas9 based approach (**S1 Fig A, B**) to knock-out two key components of mTORC2 in dHL60 cells - its scaffolding sub-unit Rictor and its kinase accessory subunit mSin1 (**Fig 1B, C**). Rictor null (Rictor KO) cells impair both mTORC2 kinase and non-kinase roles, whereas mSin1 null (mSin1 KO) cells specifically affect the kinase roles (Guertin et al., 2006; Jacinto et al., 2006, 2004; Sarbassov et al., 2004). As a pharmacological approach to impair the kinase function of mTOR, we used a specific inhibitor of mTOR Kinase Ku-0063794 (KU) (García-Martínez et al., 2009). To read out the kinase activity of mTORC2, we assayed the phosphorylation of the well-characterized mTORC2 substrate Akt (Ebner et al., 2017a; Sarbassov et al., 2005). When stimulated by chemoattractant peptide formyl-Met-Leu-Phe (fMLP); neutrophil-like differentiated HL60 (dHL60) cells derived from both Rictor KO and mSin1KO lines show marked reduction in the phosphorylation of Akt to levels that were comparable to pharmacological inhibition of the mTOR kinase (∼ 75-80% reduction from wildtype cells; **S1 Fig C, D**), indicating a loss of mTORC2 kinase activity with all of these perturbations.

To determine the importance of these mTORC2 perturbations on cell movement, we performed a transwell chemotaxis assay (**Fig 1D**), which demonstrated that the perturbation of mTORC2 complex formation (in Rictor KO) or kinase activity (in Rictor KO, mSin1KO or mTOR kinase inhibition) all led to significant and comparable defects in bulk transwell migration (**Fig 1D**; 60-70% drop in migration index compared to WT). These results agree with earlier reports of chemotaxis defects in Rictor shRNA knockdowns and upon long term perturbation of mTOR activity with Rapamycin (Diz-Muñoz et al., 2016; He et al., 2013; Liu et al., 2010). In addition, we find that both acute (with drug KU) and chronic perturbation (mSin1 KO) of mTOR kinase also impair transwell migration. This is consistent with chemotaxis defects observed in genetic knockouts of Rip3 (Sin1 ortholog of mSin1) in *Dictyostelium* (Lee et al., 2005) but is in contrast with an earlier study using partial knockdown of mSin1 in neutrophils, which found no discernible defects in chemotactic movement of mSin1 KD cells in a micro-needle assay (He et al., 2013). Factors like differences in the extent of depletion and the type of migration assay conditions used can often lead to confounding results, so we chose to assess motility defects in cleaner single cell assays of cell migration with the genetic nulls of mTORC2 components.

We suspected that mTORC2-based mechanoadaptation might be particularly important when cells are in migration environments that mechanically perturb them, including squeezing through a pore for transwell assays or migrating under mild confinement. To study 2D migration under mild confinement, we made use of under-agarose (2% w/v) overlay on cells (Bell et al., 2018; Brunetti et al., 2022; Tsai et al., 2019) and tracked individual cells over the course of 12 min (assay schematic and cell tracks in **Fig 1E**) in presence of uniform chemoattractant (25 nM fMLP). Both Rictor KO and mSin1 KO cells exhibited less net displacement (**Fig 1E**) and moved at significantly slower speeds (**Fig 1F**; median speed for Rictor KO: 3.2 μm/min; mSin1 KO: 6 μm/min) compared to Wildtype (WT) dHL60 cells (median speed: 9 μm/min). However, under similar assay environment, only Rictor KO cells showed a significantly reduced persistence (ratio of displacement/distance; median persistence for Rictor KO: 0.1; WT: 0.5), suggesting a role for Rictor beyond the kinase activity of mTORC2. In contrast to the defects observed under agarose, when Rictor KO and mSin1 KO cells were assayed for classical unconfined 2D-motility (**S1 Fig C**) on glass coverslips coated with fibronectin, they moved with similar speed and persistence as WT dHL60 cells (**S1 Fig D, E**). These results suggest that migratory defects upon mTORC2 perturbation are sensitised in an assay where cells need to actively assess the local environment and adapt during movement. These results also indicate that kinase activity of mTORC2 is specifically required to set the speed of movement, while the scaffolding roles of the complex contribute to maintain persistence of motion (**Fig 1H**).

### Kinase-independent roles of Rictor/mTORC2 restricts the zone of F-actin assembly to the cell front

Persistent motility in neutrophils relies on establishing a single front of lamellipodial F-actin. Earlier studies have shown fMLP stimulated neutrophils with reduced levels of Rictor (via shRNA knockdowns) show elevated steady-state levels of F-actin (Diz-Muñoz et al., 2016) with a near-uniform peripheral distribution of actin polymerization (Liu et al., 2010). While this is consistent with Rictor/mTORC2 mediating negative feedback to inhibit F-actin polymerisation, the relative contributions of mTORC2 kinase activity and kinase-independent roles in this process remains unclear (**Fig 2A**). Answering these questions necessitates the larger suite of mTORC2 perturbations that we leverage in the current work.

**Figure 2.**
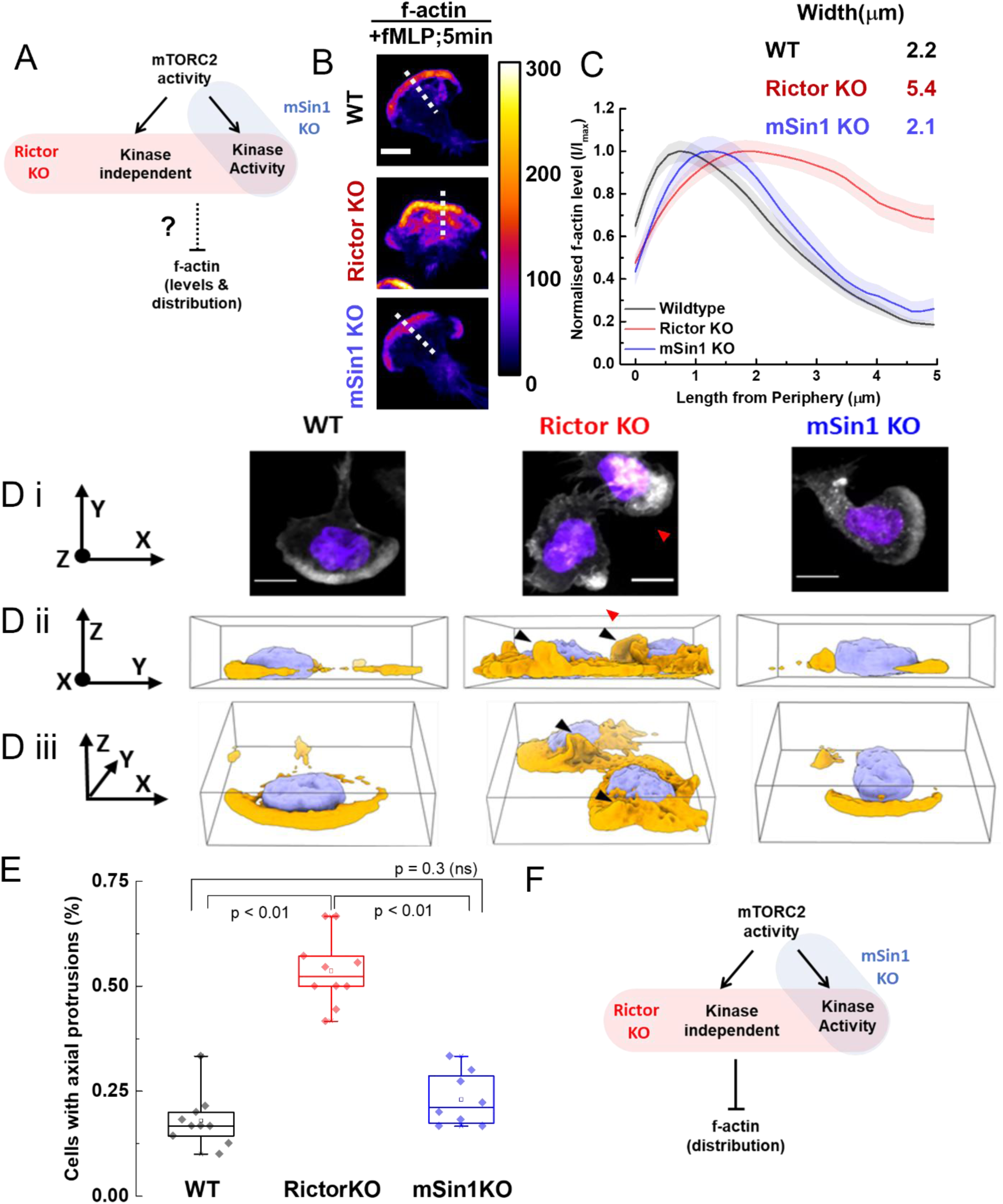
Kinase-independent arm of Rictor/mTORC2 restricts the zone of f-actin assembly to the front of the cells. (A) Schematic shows how we are probing the kinase-dependent vs independent roles of mTORC2 in regulating f-actin levels and spatial organisation. (B) Maximum intensity projections of Alexa647-phalloidin stained F-actin obtained from confocal z-stacks (10 μm deep) for wildtype (WT), Rictor KO, or mSin1KO dHL60 cells, 5min after stimulation with 25nM fMLP. Fire-LUT shows the intensity scaling; scale bar 10 μm. (C) F-actin intensity (normalised to individual peak) line-scans (Mean ± SEM) obtained (dashed lines on B) from wildtype (n=35), Rictor KO (n=48), or mSin1KO (n=26) dHL60 cells obtained from two independent experiments. Rictor KO (red) have a wider lateral zone of leading edge F-actin in comparison to wildtype (WT; black) and mSin1KO (blue); quantified by Bi-gaussian fitting of the intensity profile. (D) Representative wildtype (WT), Rictor KO and mSin1KO dHL60 cells shown as either maximum intensity projection (in *xy-plane*; D I; scale bar 10 μm); or a ChimeraX 3D-reconstruction in *yz-plane* (D ii) and a tilted *xz-plane* (D iii) to highlight the axial features of F-actin distribution. Rictor KO cell shows protrusions out of the plane of the substrate that are rarely present in either Wildtype (WT) or mSin1KO cells. (E) Box-plots quantify fraction of cells with axial protrusions obtained from ChimeraX 3D-reconstructions views of each cell type (∼10 fields; at least 100 cells analysed for each condition) across two independent experiments. RictorKO cells have significantly (p < 0.01, one-way ANOVA with Tukey’s mean comparison test) higher fraction of cells with axial protrusions. For box plots, median is indicated by the line, inter-quartile range sets the box width and error bars indicate 10-90^th^ percentile. (F) Defects observed in f-actin distribution for only RictorKO dHL60 cells but not mSin1KO cells suggests kinase independent arm of Rictor/mTORC2 is required for negative feedback on f-actin assembly, distribution, and organisation.

To assay actin assembly in our mutant backgrounds, we stimulated fibronectin-adhered wildtype (WT), Rictor KO and mSin1 KO dHL60 neutrophils with chemoattractant and stained them with Phalloidin-Alexa647. WT cells show an initial burst (1 min, **S2 Fig A, B**) in actin polymerisation that adapts over 5 minutes to basal (0 min, no fMLP) levels. Consistent with our earlier results from partial loss of Rictor (in knockdown cells), Rictor KO cells exhibit significant increase in F-actin levels (1 min ; **S2 Fig A, B**). In contrast, depletion of mSin1 failed to elicit appreciable differences in the overall levels during the initial burst and reset (mSin1 KO, **S2 Fig A, B**), suggesting the kinase arm of mTORC2 may be dispensable for regulating F-actin levels. To investigate the role of mTORC2 on the spatial dynamics of actin regulation, we next focused on the subcellular features of F-actin distribution in polarised cells after 5 minutes of chemoattractant stimulation (**Fig 2B**), visualized via maximum-intensity projections of 3D data stacks. Rictor KO cells showed broader range of F-actin in the front compared to WT and mSin1 KO cells (**Fig 2B**). We used linescan (5 μm line ROI) to measure F-actin levels orthogonal to the leading front (**Fig 2C**) and fitted the F-actin intensity profile to a Bi-Gaussian to estimate the effective width of the F-actin zone (Buys and De Clerk, 1972). While WT and mSin1KO cells have very similar widths of F-actin zone (fitted width ∼ 2 - 2.5 μm), Rictor depletion led to a two- fold expansion of the width of the actin front (fitted width 5.4 μm).

Recent lattice light sheet imaging data have shown neutrophils not only generate substrate-bound protrusions but also build axial protrusions that extend away from the plane of substrate (Fritz-Laylin et al., 2017; Pipathsouk et al., 2021). To investigate whether Rictor/mTORC2 plays a role in constraining the formation and abundance of these axial protrusions, we analyzed the extent of protrusion formation in 3D reconstructions of neutrophils. We used ChimeraX (Pettersen et al., 2021) to 3D reconstruct and render these cells in two axial planes *yz* and *tilted xz* (**Fig 2D ii, iii, Video 1**). Confirming our expectation, loss of Rictor led to enhanced accumulation of F-actin rich protrusions away from the plane of the substrate. These protrusions were frequently present (> 50 % of all cells imaged; **Fig 2E, Video 1**) in Rictor KO cells, and they were more rarely observed in WT or mSin1 KO cells (∼ 25% of all cells; **Fig 2E**).

The absence of discernible defects in F-actin distribution in absence of mSin1 (and hence mTORC2 kinase activity) suggest that mTORC2 relies on its Rictor-dependent kinase independent signaling to restrict F-actin to the leading edge of cells. Next, we investigated how mTORC2 regulates the biochemical effectors of front/back polarity in cell motility.

### Kinase-independent arm of Rictor/mTORC2 inhibits Rac activity while its kinase role stimulates myosin contractility

The migration phenotypes we observe for both kinase-dependent and kinase-independent arms of Rictor/mTORC2 (**Fig. 1**) could arise from its effects on different portions of migration cascade. A wide range of motile cells including neutrophils show a distinct front-back polarity and organise their protrusive fronts and contractile backs using Rac and RhoA/myosin signaling, respectively (Ku et al., 2012; Schaks et al., 2019; Xu et al., 2003). We were interested in how mTORC2 regulates these polarity and cytoskeletal programs. Cellular stretch has been shown to inhibit Rac activity in several contexts (Houk et al., 2012; Katsumi et al., 2002), but it is not clear which arm of mTORC2 activities are necessary for this inhibition in neutrophils.

Here, we verified whether kinase activity of mTORC2 is dispensable restricting Rac activation and actin polymerization (**Fig 3A left side**). To investigate mTORC2 regulation of Rac, we leveraged a biochemical assay of Rac activity following the phosphorylation profile of Rac effector p21 kinase (Pak) upon chemoattractant stimulation (Graziano et al., 2019; Weiner et al., 2006). Since chemoattractant (fMLP) addition is sufficient to trigger downstream signaling and polarisation of cells in suspension, we carried out these biochemical assays in suspension. In WT dHL60 neutrophil-like cells (**Fig 3B, C**), fMLP addition led to a burst of Pak-phosphorylation (Rac activity) that peaks around 1min followed by gradual adaptation over the course of 5 minutes. In comparison, Rictor KO cells exhibit significantly elevated levels (∼ 2 fold above WT; 1-2 min; **Fig 3C**) of Rac activity upon fMLP addition. However, perturbation of mTORC2 kinase activity (in mSin1 KO cells) did not result in appreciable alteration in the temporal profile of Rac activity. The marked differences in Rac activity observed upon depletion of Rictor or mSin1 suggests that kinase-independent roles of Rictor/mTORC2 are responsible for limiting leading edge signals.

**Figure 3.**
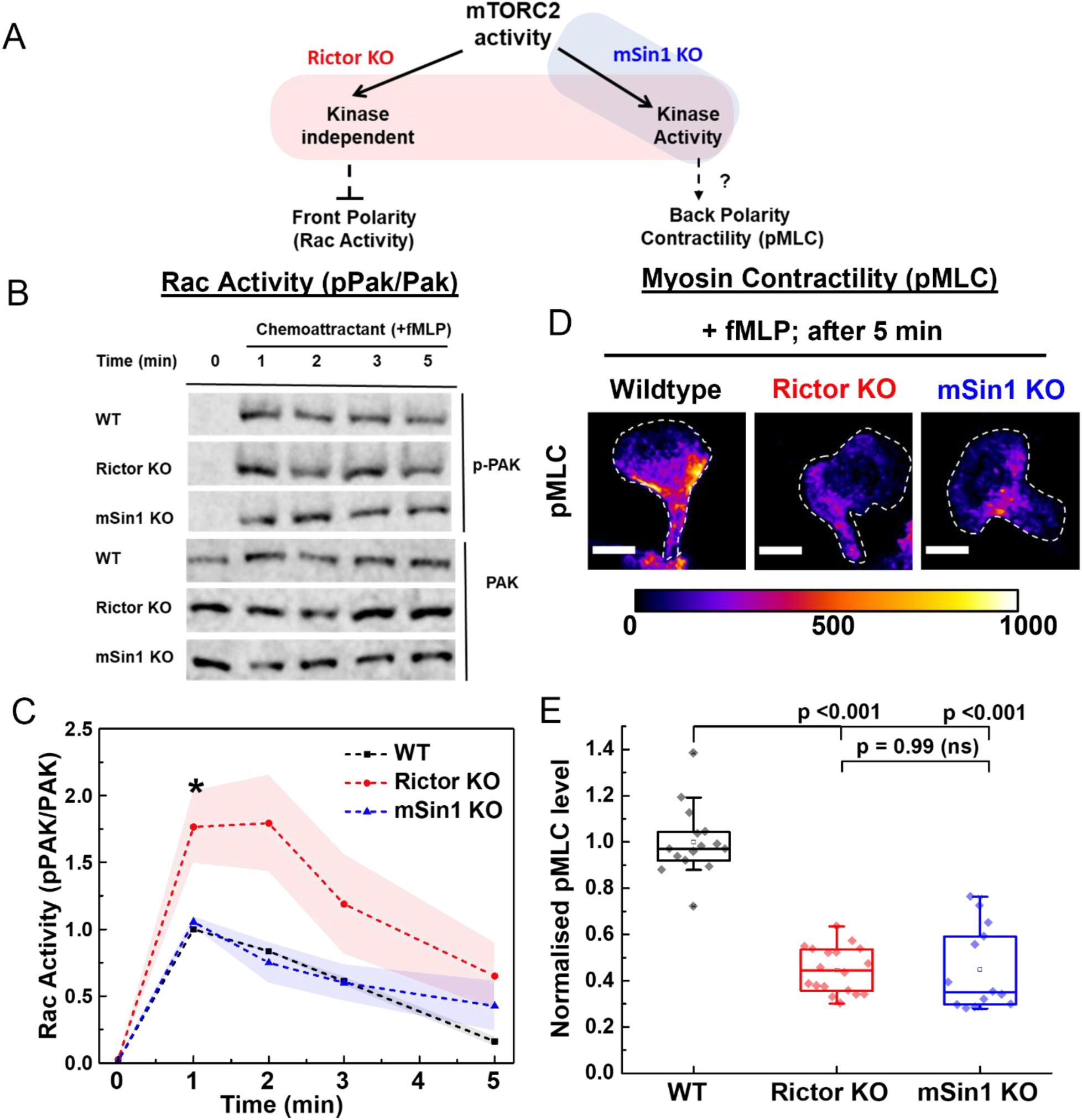
Kinase-independent arm of Rictor/mTORC2 inhibits Rac activity while its kinase role stimulates myosin contractility. (A) Schematic shows how we probe the kinase-dependent vs independent roles of mTORC2 in regulating front (Rac/f-actin) and back polarity (RhoA/myosin) programs. (B) Rac activity was quantified for chemoattractant (25nM fMLP) stimulated dHL60 cells (wildtype, RictorKO and mSin1KO) using antibodies targeting phospho-PAK (pPAK), a downstream readout of Rac activation. Antibodies against total PAK were used as a loading control and calculate the ratio of pPAK/PAK, the readout for Rac Activity we use here. (C) Rac-activity (assayed by pPAK/PAK) of wildtype(WT), RictorKO, or mSinKO cells upon stimulation with 25nM fMLP; each plot shows Mean ± SEM from three-independent experiments, with 1- min timepoint of wildtype (WT) being used to normalise all conditions for each independent trial. * p<0.05 by unpaired t-test between Rictor KO and WT at 1 min timepoint. (D) Phospho-myosin light chain (pMLC) immunostaining (labelled with Alexa488 secondary antibody) of wildtype(WT), Rictor KO and mSin1KO cells, 5 mins after stimulation with 25nM fMLP. Images show maximum intensity projections obtained from 10 μm deep confocal z-stacks; scale bar 10 μm. Dashed outlines indicate the cell boundary, and all conditions were equally intensity scaled as shown by associated Fire LUT. (E) Box-plots show the quantification of total pMLC intensity levels from confocal z-stacks as shown above (∼ 15 fields and at least 150 cells; pooled from 2 independent experiments) across each condition for the three cell types. Both mutant cell types have significantly diminished pMLC levels compared to wildtype cells (p < 0.001, one-way ANOVA with Tukey’s mean comparison test), suggesting that the kinase activity of Rictor/mTORC2 stimulates myosin contractility. For box plots, median is indicated by the line, inter-quartile range sets the box width, and error bars indicate 10-90^th^ percentile.

Next, we investigated whether perturbation of mTORC2 activity also affects polarity signaling at the trailing edge of the cell. Active RhoA and its associated RhoA-Kinase (ROCK) localize and phosphorylate myosin regulatory light chain (pMLC) to power contractile retraction of the back (Hind et al., 2016; Tsai et al., 2019; Wong et al., 2006). Biochemical studies in neutrophils have shown that mTORC2 kinase effector PKC regulates RhoA and myosin activity in neutrophils (Liu et al., 2014, 2010). In fission yeast, TORC2 kinase activity also stimulates myosin activation (Baker et al., 2019) indicating a conserved role for TORC2 kinase effectors in regulating contractility. If kinase roles of mTORC2 stimulate myosin contractility, we expect both Rictor KO and mSin1 KO cells to show reduced pMLC levels (Schematic in Fig 3A). If the Rho/myosin inhibition is secondary to an increase in the antagonistic Rac activation, we would expect larger effects for Rictor KO cells. (**Fig 3A right side**). We used immunofluorescence (IF) against phospho-MLC to probe myosin contractility in dHL60 neutrophils plated on glass and stimulated with chemoattractant (25 nM fMLP). Wildtype neutrophils (S3 Fig A) gradually elevate their pMLC levels (and hence contractility) over the course of 5 minutes as they polarise (**Fig 3D, S3 Fig B**). While Rictor KO and mSin1 KO cells have comparable levels of basal pMLC (0 min) as wildtype; both cell types fail to raise their pMLC levels (∼ 50-60 % reduction) over the course of polarisation (**Fig 3D, S3 Fig A, B**). We found the contractility loss upon mTORC2 perturbation were comparable with direct inhibition of RhoA/ROCK activity. Wildtype cells treated with ROCK inhibitor Y27632 and stimulated with fMLP (5 minutes) show significant loss (∼80% drop) of pMLC levels and emergence of a long stalk (**S3 Fig C**) as cells fail to retract. In comparison, pMLC levels remain unchanged between untreated and Y27632 treated Rictor KO cells (∼60 % less than WT ; **S3 Fig D**).

Our results show that the kinase-independent roles of Rictor/mTORC2 inhibit Rac activity in the front, while the kinase roles of mTORC2 stimulate contractility at the back allowing the two divergent downstream arms of mTORC2 to execute opposing effects on the front and back polarity programs. We next explore the consequence of this regulatory circuit for front-back polarity coordination during motility.

### Rictor/mTORC2 spatially and temporally coordinates the front and back polarity program during movement

Migrating neutrophils rely on front-back coordination to persistently move, turn, and reorient during migration. This front-back coordination depends on multiple currencies including cell membrane tension (Houk et al., 2012; Sens and Plastino, 2015), actin flows (Liu et al., 2015; Maiuri et al., 2015), myosin contractility (Tsai et al., 2019), as well as biochemical signaling crosstalk (Devreotes et al., 2017; Ku et al., 2012; Xu et al., 2003) between the front and back polarity programs. Our results (**Fig 3**) suggest a scenario in which neutrophils appear to leverage the two arms of mTORC2 downstream signaling to coordinate front and back polarity signaling. We asked whether this logic contributes to the spatial organisation and temporal coordination of polarity during movement (**Fig 4**).

**Figure 4.**
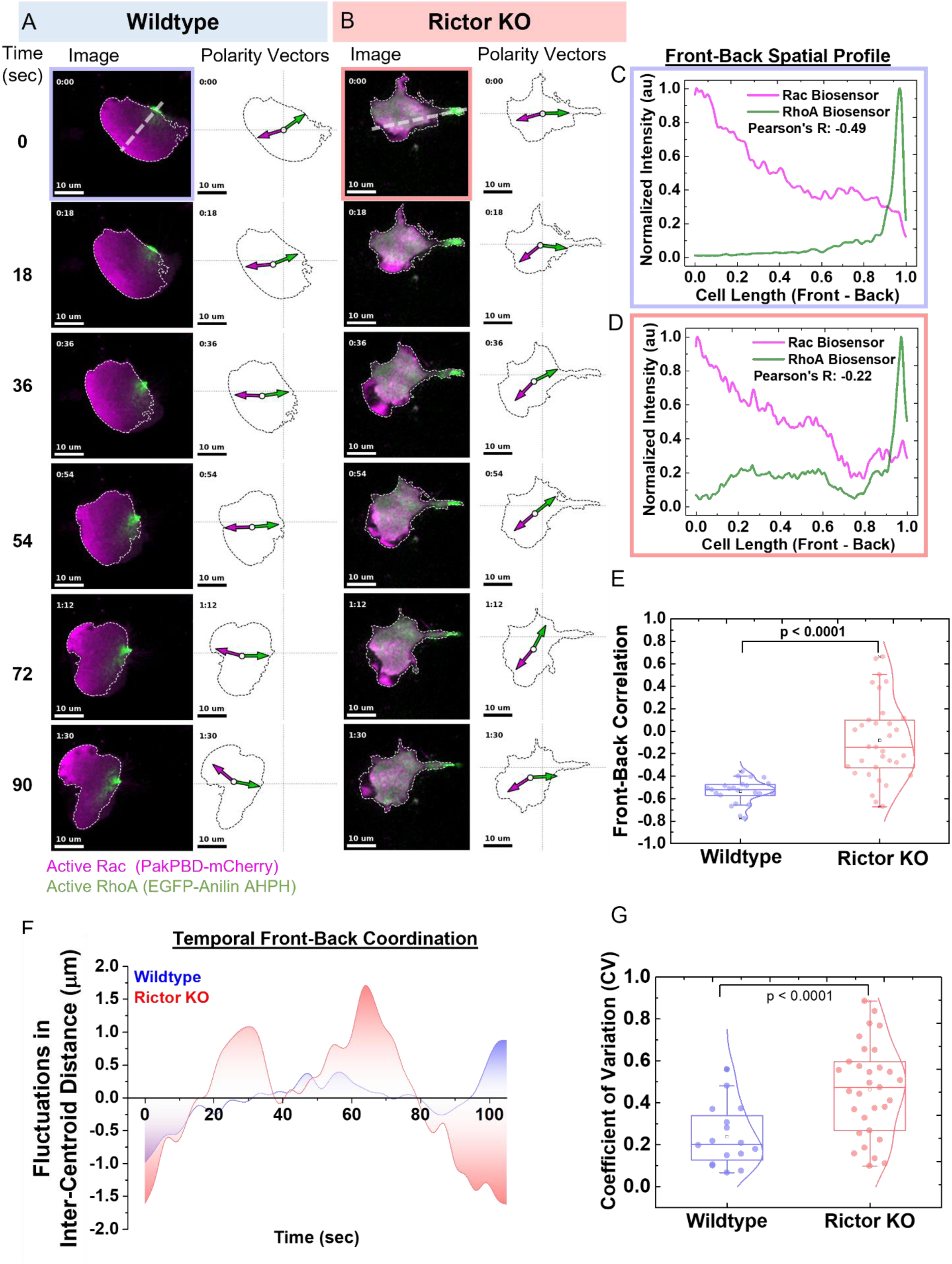
Rictor/mTORC2 is required for maintaining the spatial and temporal coordination of the front and back polarity programs. (A) Wildtype (WT) and Rictor KO (B) cells co-expressing Rac biosensor (Pak-PBD-mCherry) and RhoA biosensor (EGFP-Anillin-AHPH) were plated under 2% agarose overlay and imaged every 3 seconds using a confocal spinning disk microscope. Montage of images acquired over 90 sec show the distribution of front (Rac*, magenta) and back (Rho*, green) polarity signals (Image, left) and the corresponding front (magenta arrow) and back (green arrow) polarity vectors. The cell centroid for each frame is indicated by the open circle and it’s displacement from the grid indicates overall cell movement; scale bar is 10 μm. (C, D) Anti-correlation between the intensity profile of polarity signals across the front-back axis provides a measure of spatial segregation of the front-back signals. Representative intensity profiles of Rac* and RhoA* obtained from line-scan (dashed line in time 0 sec in A, B) of wildtype (WT, C) and Rictor KO (D), with computed Pearson’s correlation coefficient for each set of intensity traces (Pearson’s correlation coefficient = - 0.49 for WT; -0.22 for RictorKO). More strongly negative Pearson’s correlation coefficient indicates better separation between front and back signals. (E) Box-plots of front-back correlation values for wildtype (WT, n=23 cells) and Rictor KO cells (n= 33 cells) pooled from 4 independent experiments. Rictor KO cells have significantly higher correlation coefficient (p<0.0001; Mann-Whitney’s test) compared to wildtype suggesting impaired spatial sorting of front-back polarity programs. (F) To measure the extent of temporal coordination between polarity signals, we analyzed the fluctuations in the weighted inter-centroid distance between the front and back polarity biosensor intensities (indicated by the distance between the arrowheads of polarity vector in images A, B). Wildtype cells have polarity vectors uniformly aligned to the front-back axis and maintain a stable inter-centroid distance (blue), while RictorKO cells show stronger fluctuations in inter-centroid distance (red). Representative plot of fluctuations in inter-centroid distance for both cell types of WT and Rictor KO. We use coefficient of variation (CV) as a metric to quantify the magnitude of the fluctuations. (G) Box-plots of distribution of CV obtained for wildtype(n =19 cells) and RictorKO (n =30 cells) across four independent experiments. Rictor KO cells exhibit significantly higher fluctuations in ICD(p=0.0006; Mann-Whitney’s test), suggesting impaired temporal coordination of front and back polarity programs in these cells. For box plots, median is indicated by the line, inter-quartile range sets the box width, and error bars indicate 10-90^th^ percentile.

To investigate the role of mTORC2 in front/back coordination, we revisited the agarose overlay conditions (Bell et al., 2018; Brunetti et al., 2022) which sensitized Rictor-dependent migratory phenotypes (Fig 1E-G). To map the distribution and coordination of polarity in migrating cells, we co-expressed localization biosensors for both active Rac (from Pak; Srinivasan et al., 2003) and active RhoA (from Anilin; Piekny and Glotzer, 2008) in wildtype and Rictor KO cells and imaged them at high spatial and temporal resolution (frame interval of 3 second) in presence of uniform chemoattractant 25 nM fMLP (**Fig 4A, B montage, Video 2**). First, we assessed the spatial features of front and back regulation in these cells. Active Rac forms a gradient in front half of a polarised wildtype cell, while active RhoA is tightly focused at the rear (**Fig 4A**, WT). A linescan in the direction of cell movement (front-back axis) reveals the mutually exclusive distribution of biosensor and a strong anti-correlation (Pearson’s R = 0.49 for WT; **Fig 4C**). While Rictor KO cells also polarise, they do so with an elongated stalk at the cell back (**Fig 4B**, Rictor KO), where the bulk of active RhoA is concentrated. As expected, wildtype cells showed strong mutual antagonism of Rac and RhoA, with RhoA activity completely excluded from the Rac-positive zones in the cell front (**Fig 4C**). In contrast, Rictor KO cells showed local hotspots of RhoA biosensor localisation in the cell front (**Fig 4D**). This impaired spatial sorting of front and back polarity signal can be detected by poor anti-correlation between both biosensors (Pearson’s R = -0.22 for Rictor KO; **Fig 4D**). Front-back correlation (**Fig 4E**) quantified from several cells (WT = 23 cells; Rictor KO = 33 cells) show a significant difference between the two cell types and overall shift in the distribution to poorer front/back separation for Rictor KO (Median for WT= - 0.55; Rictor KO = 0.1). These defects originate specifically from differences in biosensor distribution across the front-back axis, as lateral linescan perpendicular to direction of movement (**S4 Fig A, B**) do not show any preferential bias and resulted in similar overall distribution of Pearson correlation across the lateral plane for both wildtype and Rictor null cells (**S4 Fig C**).

Next, we probed the temporal coordination of front-back polarity during movement under agarose. Recently, we and others have used a polarity metric based on the centroid of biosensor intensity with respect to the geometric centroid of the cells to monitor Rac polarity (Graziano et al., 2019; Olguin-Olguin et al., 2021). We revised this analysis approach to compute the centroid of biosensor intensities for both the Rac and RhoA biosensor (polarity vectors for WT and Rictor KO in **Fig 4A, B**). The distance between the zones of Rac and RhoA activation (inter-centroid distance) provides a measure of separation between front and back signals during migration (length between arrowheads of polarity vectors for WT and Rictor KO in **Fig 4A, B**). Cells proficient in coordinating front-back polarity during persistent movement exhibit smaller fluctuations in inter-centroid distance. Rictor KO cells show enhanced amplitude of fluctuations for front/back separation (**Fig 4F**). We quantified the strength of these fluctuations as coefficient of variation (CV), a commonly used metric to quantify fluctuations in cell polarity (Lai et al., 2018; Onwubiko et al., 2019). Loss of Rictor leads to a significant increase in CV (**Fig 4G**; Median CV for WT: 0.2, Median CV for Rictor KO : 0.5), indicating that Rictor/mTORC2 plays a critical role in coordination between the polarity signals at front and back.

Our experiments probe how different branches downstream of mTORC2 activation (kinase dependent vs independent) relate to regulation of front-back polarity programs and their spatiotemporal coordination during movement. We next investigated whether biochemical signals and physical forces synergize to active mTORC2 in the first place.

### Mechanical stretch synergizes with PIP_3_ generation to activate mTORC2

TORC2 is an ancient program that has been shown to be regulated by both biochemical and mechanical inputs in several contexts ranging from yeasts to Dictyostelium to immune cells. While mechanical stretch is one important regulator of the activation of the complex in *Dictyostelium* (Artemenko et al., 2016) mounting evidence suggest biochemical inputs from Ras(and Rap) are required for full activation of the complex (Khanna et al., 2016; Senoo et al., 2019; Smith et al., 2020) .

To investigate whether mechanical stretch suffices to activate the mTORC2 complex, we exposed neutrophils to hypotonic media (50% decrease in ionic strength) and assayed the activation of mTORC2 via Akt phosphorylation. Surprisingly, hypotonic exposure (purely mechanical input) alone failed to activate mTORC2, but hypotonic exposure synergized with chemoattractant addition (combination of biochemical and mechanical inputs) to activate mTORC2 (**Fig 5A**). Hypotonic buffer (50% water) failed to stimulate mTORC2 kinase activity at both 1 and 3 minutes after osmotic challenge (**Fig 5B, C**). However, a combination of osmotic challenge and chemoattractant, significantly amplifies (∼ 2 fold) the peak pAkt response at 3 minutes compared to the levels obtained by chemoattractant alone (**Fig 5C**). These results suggest that mechanical stretch synergizes with other biochemical inputs from chemoattractant to amplify mTORC2 signaling activity.

**Figure 5.**
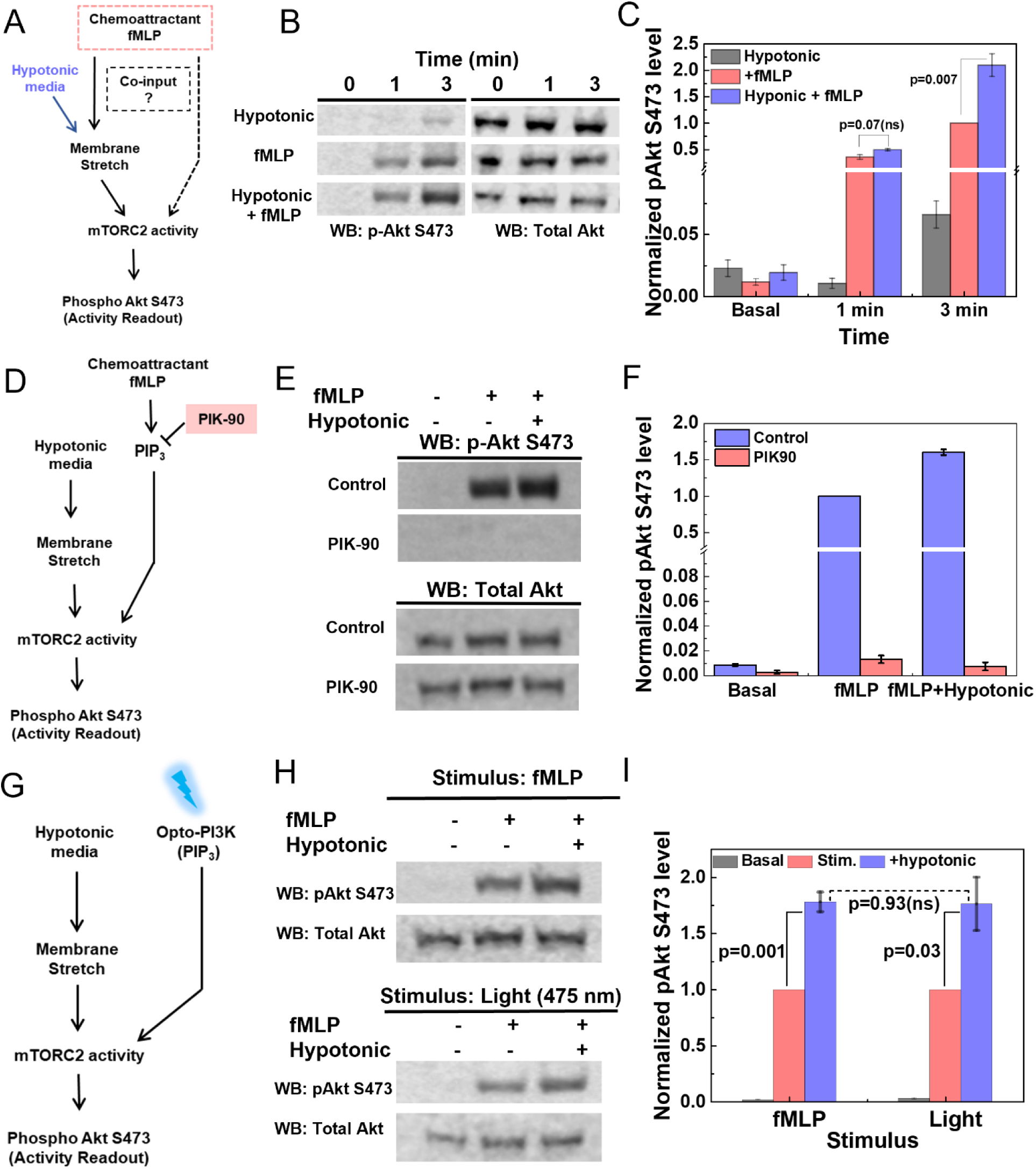
Mechanical stretch synergizes with PIP3 generation to activate mTORC2. (A) Probing whether mTORC2 activation can be mediated by mechanical stretch alone (here simulated by hypo-osmotic shock) or whether it requires additional biochemical inputs downstream of chemoattractant stimulation. (B) To probe the logic of mTORC2 activation, dHL-60 cells were subjected to either just hypotonic media (50% osmolarity reduction), stimulated with 20nM fMLP (fMLP only), or subjected to both inputs (hypotonic + 20nM fMLP). mTORC2 activity was assayed using phospho- Akt/pan-Akt immunoblots, and representative immunoblot panels are shown. (C) mTORC2 activity was quantified and normalised using fMLP (3 min) data for each experiment. Hypotonic shock (grey bars) alone doesn’t stimulate mTORC2 activity and chemoattractant addition is needed to trigger activation of signaling (red bar). In the presence of fMLP, hypotonic shock (blue bar) amplifies the signaling output of mTORC2 (p = 0.007, unpaired t-test. Each bar reflects Mean ± SEM from three independent experiments. (D) Testing whether PIP_3_ is necessary to stimulate mTORC2 activity with the PI3Kγ inhibitor PIK-90. (E) Representative immunoblots of pAkt and Akt to measure mTORC2 activation. Control (top) or PIK- 90 (1 μM; bottom) treated dHL60 cells were assayed for mTORC2 activity in absence (basal activity; -/- condition; left lane) or in presence of fMLP alone (+/-; middle lane) or a combination of fMLP and hypotonic shock (+/+; right lane). In presence of PIK-90, mTORC2 activity is severely attenuated (with background levels of pAkt detected). (F) mTORC2 activity quantified across the three conditions and normalised using the fMLP (Control) data for each experiment. Each bar reflects the mean ± SEM obtained from three independent experiments. Inhibition of PI3K activity shows PIP_3_ is a necessary co- input for activation (fMLP) and amplification of mTORC2 activity upon stretching (fMLP + hypotonic). (G) Testing whether PIP_3_ (via optogenetic stimulation) is sufficient to activate mTORC2 and can collaborate with mechanical stretch for mTORC2 activation. (H) Representative immunoblots of pAkt and AKT for dHL60 cells stimulated with either chemoattractant (20 nM fMLP for 3 min; top panel) or light at 475 nm (1mW for 3 min) to stimulate Opto-PI3K (bottom panel). For both inputs, pAkt/AKT was assayed for basal (left lane; -/-), just stimulus (middle lane; +/-) or when paired with hypotonic media (right lane; +/+). (I) mTORC2 activity (assayed by pAkt/AKT ratio) quantified across the three different conditions (H) for both stimulus (fMLP or light) and normalised using the stimulus only condition (just fMLP or light) for each experiment. Activation of mTORC2 and amplification of signaling activity upon hypotonic shock (p =0.001 for fMLP; p =0.03 for Light; both by unpaired t-test) show similar behavior (p = 0.93, ns, unpaired t-test) when either fMLP or opto-PI3K was used as stimulus. Each bar reflects mean ± SEM for 3 independent experiments.

Chemoattractant fMLP activates a wide range of downstream signaling pathways (Gβγ, PI3K, Ras; Xu et al., 2003). Which of these programs are necessary to synergize with mechanical inputs to stimulate mTORC2? Here we focused on the PI3K node that is responsible for PI-3,4,5-P3 (PIP_3_) generation at the front of the cells. PIP_3_-Rac positive feedback is central for instructing actin polymerisation in the front and raising cellular tension (Graziano et al., 2017; Wang et al., 2002; Weiner et al., 2002). Moreover, PIP_3_ is also responsible for recruitment of mTORC2 phospho-substrate Akt (Ebner et al., 2017b, 2017a). In neutrophils, PI3Kγ is the dominant regulator of fMLP-dependent PIP_3_ production (Hannigan et al., 2002; Hirsch et al., 2000; Sasaki et al., 2000; Stephens et al., 1994). To test if PI3Kγ activity is a necessary for stretch-dependent amplification of mTORC2 activity we used PIK90 (Van Keymeulen et al., 2006), a specific inhibitor of PI3Kγ (**Fig 5D**). dHL60 cells pre-treated with PIK90 and subsequently stimulated with fMLP alone or in combination with hypotonic media (50 % ionic strength) show background (basal) levels of pAkt in immunoblots compared to untreated cells (**Fig 5E, F**). These results show that PI3Kγ induced PIP_3_ synthesis is necessary for neutrophils to activate and amplify mTORC2 kinase activity upon osmotic stretch.

We next tested whether PIP_3_ suffices to replace chemoattractant in mechanics-synergized mTORC2 activation. For this purpose, we used an optogenetic module (opto-PI3K) which can synthesize PIP_3_ in response to light stimulation (Graziano et al., 2017) (**Fig 5G**). Optogenetic stimulation of PIP_3_ sufficed as the co-input with mechanical stretch in mTORC2 activation (**Fig 5H, I**). Our results indicate that mTORC2 is jointly activated by PIP_3_ and mechanical stretch.

## DISCUSSION

For persistent motility, neutrophils must coordinate the activation of their leading and trailing cytoskeletal networks (Houk et al., 2012; Tsai et al., 2019). The mTORC2 complex is a critical regulator of this coordination. Following protrusion-induced membrane stretch, mTORC2/Rictor activation inhibits actin polymerization to enable a dominant front to emerge (Diz-Muñoz et al., 2016) and regulates myosin contractility at the trailing edge (Liu et al., 2010). Here we use a combination of genetic nulls and pharmacological tools to investigate the kinase dependent and independent links from mTORC2 to these cytoskeletal programs. mTORC2 is indispensable for movement when neutrophil-like dHL60 cells are mildly confined (∼ 5 μm) under agarose to mimic the confined spaces these cells explore *in vivo* (**Fig 1**). The kinase-independent roles of Rictor/mTORC2 are central to regulating the leading edge polarity program (Rac activity, F-actin distribution; **Fig 2 & 3**). Consistent with earlier studies (Baker et al., 2019; Liu et al., 2014, 2010), we find that mTORC2 kinase activity is essential for sustained myosin contractility at the trailing edge (**Fig 3**). Using dual biosensor imaging of front and back polarity program, we show that mTORC2 is necessary for these two arms to work in unison for spatial and temporal coordination of polarity during movement (**Fig 4**). Finally, we probe the requirements for mTORC2 activation. Membrane stretch does not suffice for mTORC2 activation unless the biochemical input PIP_3_ is also present (**Fig 5**). In summary, our results highlight the logic of stretch-activated Rictor/mTORC2 signaling in coordinating front-back polarity during neutrophil movement (working model in **Fig 6**).

**Figure 6.**
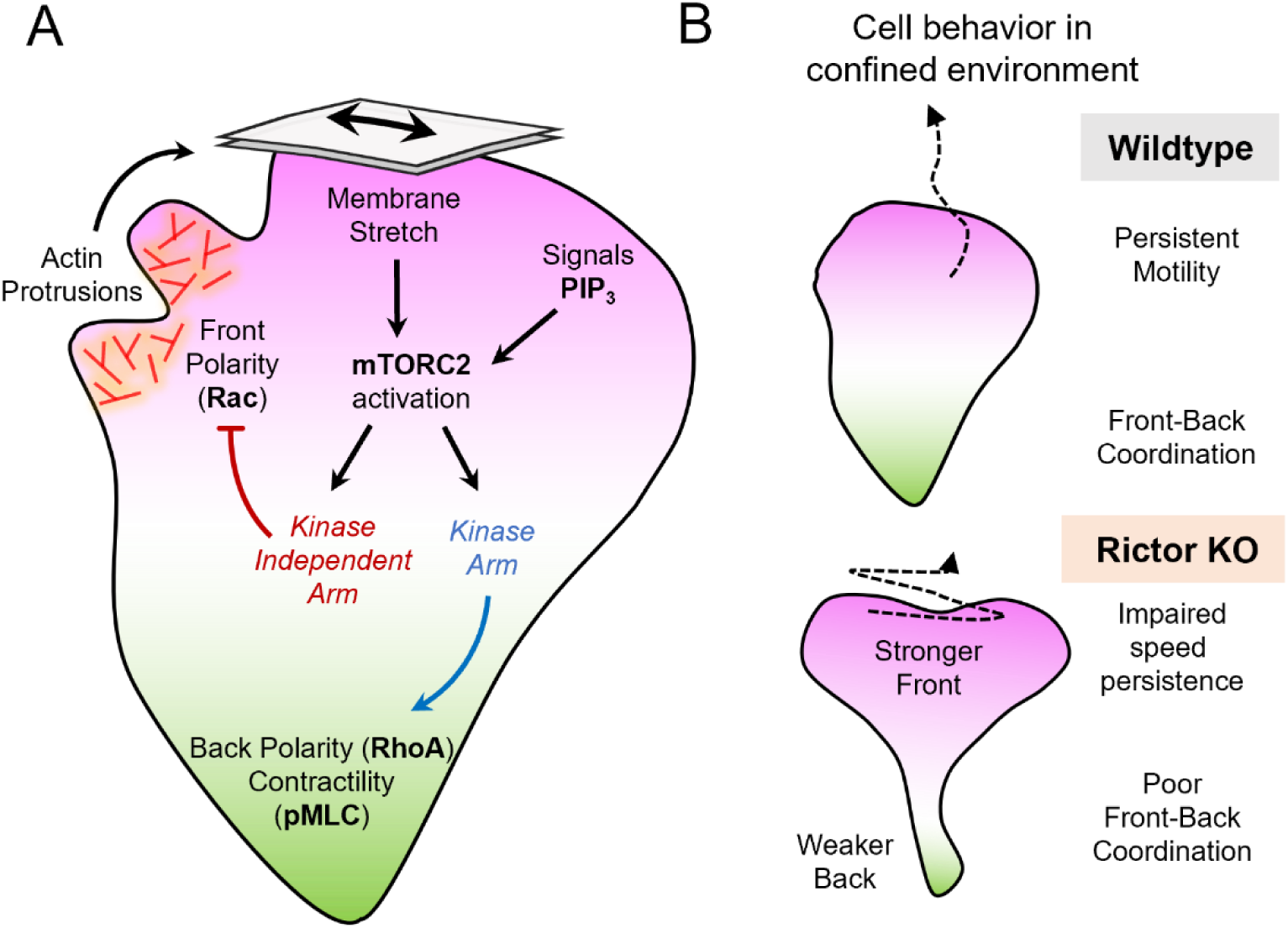
Working model for the molecular logic of mTORC2 based regulation of front and back cytoskeletal programs. (A) Membrane stretches and biochemical signals from PIP3 synergize to activate kinase and kinase independent roles of mTORC2. Non-catalytic kinase independent roles of Rictor/mTORC2 allow stretch-dependent inhibition of front polarity signals (Rac) and restrict F-actin protrusion to the leading edge. The kinase roles of mTORC2 stimulate myosin contractility (pMLC) at the back. This bifurcation of the downstream mTORC2 activities enables independent regulation of the spatially polarised front (magenta) and back (green) programs and coordination between them. (B) The regulatory circuit for mTORC2 based front-back coordination (A) is essential for persistent movement in confined environments where cells experience mechanical stretch (like under agarose). In absence of mTORC2 activities (as in Rictor KO) cells exhibit elevated Rac activity (Stronger Front) and lowered contractility (Weaker Back); consequently front-back coordination is lost resulting in impaired speed and persistence of motility in confined environment.

TORC2 is an ancient regulator of plasma membrane (PM) tension homeostasis across evolution (Eltschinger and Loewith, 2016; Riggi et al., 2020). In yeasts, an increase in PM tension triggers TORC2-based homeostatic responses to increase the surface area of the plasma membrane (Berchtold et al., 2012; Riggi et al., 2019, 2018; Roelants et al., 2017). In Dictyostelium, the TORC2 signaling program is activated by shear stress and regulates polarity, chemotaxis and electrotaxis (Gao et al., 2015; Kamimura et al., 2008; Lee et al., 2005). Membrane-tension based regulation of actin polymerization appears to be a conserved mechanism to reinforce polarity and cortical architecture (Sens and Plastino, 2015). In neutrophils, disruption of mechanosensory mTORC2 leads to elevated levels of membrane tension arising from increased actin polymerisation (Diz-Muñoz et al., 2016). Among other immune cells, loss of Rictor in B-cells also led to drastic increase in cortical F-actin levels upon B-cell receptor (BCR) stimulation (Huang et al., 2017). Similar mechanisms could also restrict actin polymerisation at the tip of the leading edge of oligodendrocytes as they wrap around the axonal shaft during myelin sheath formation (Bercury and Macklin, 2015; Nawaz et al., 2015), as mTORC2 signaling has been shown to regulate the differentiation, shape, and actin cytoskeleton organisation of these cells (Bercury et al., 2014; Dahl et al., 2022).

How does mTORC2 interface with cell polarity? While the kinase-independent roles of Rictor/mTORC2 inhibits the front (Rac); its kinase activity stimulates the back (RhoA). Using two distinct aspects of mTORC2 potentially enables independent regulation of the spatially polarised front and back program. Further, the use of a shared signalling node of mTORC2 activation may help facilitate coordination between the two polarity programs (**Fig 6**). This mTORC2-based mechanism of signaling based coordination between the two programs is likely to operate in conjunction with the recently reported fast mechanical coupling of protrusion and retraction dynamics observed in neutrophils (Tsai et al., 2019) and in other cells undergoing ameboid migration (Liu et al., 2015; Maiuri et al., 2015).

How are downstream arms of Rictor/mTORC2 linked to regulation of front/back polarity signals? We show that the kinase independent arm of Rictor/mTORC2 inhibits Rac activation and actin polymerization at the leading edge (**Figs 2, 3**) consistent with some (Diz-Muñoz et al., 2016; Liu et al., 2010) but not all (He et al., 2013) previous studies, possibly owing to differences between knockdowns in previous work and complete knockouts in the present work. There are a number of potential mechanistic links from Rictor/mTORC2 to Rac activation. mTORC2 has been shown to directly interact with Rac1 (Saci et al., 2011) and regulate Rac GEFs P-Rex1 and Tiam1 (Hernández-Negrete et al., 2007; Morrison Joly et al., 2017). Some of the kinase independent roles of mTORC2 could also arise from the scaffolding roles of Rictor independent of mTORC2 (Gkountakos et al., 2018; Smith et al., 2020). For instance, Rictor participates in mTOR independent scaffolding complex with the unconventional Myo1c in adipocytes to regulate membrane ruffing (Hagan et al., 2008), whose hematopoietic isoform Myo1g is a key regulator of cellular surface topology and membrane tension in other immune cells like T-cell and B-cell (Gérard et al., 2014; López-Ortega et al., 2016). The kinase activity of mTORC2 is essential for spatial distribution of active RhoA, sustaining myosin contractility and tail retraction (**Figs 3, 4**). This conserved role of mTORC2 kinase activity has also been reported in both fission yeast and neutrophil (Baker et al., 2019; Liu et al., 2010). While earlier studies in neutrophils had linked mTORC2 dependent phosphorylation of PKCβII to cAMP based regulation of RhoA activity (Liu et al., 2014, 2010), we further show kinase roles of mTORC2 broadly attenuates myosin contractility (via reduced pMLC) and impairs front-back coordination. Future work will identify molecular players participating downstream of mTORC2 to regulate front-back polarity signals. mTORC2 activation integrates several upstream inputs like chemoattractant, growth factors, Ras, and mechanical forces (Charest et al., 2010; Diz-Muñoz et al., 2016; Khanna et al., 2016; Kovalski et al., 2019; Smith et al., 2020). In Dictyostelium, TORC2 activation relies on inputs from other leading edge components Ras and Rap GTPase (Kamimura et al., 2008; Khanna et al., 2016; Senoo et al., 2019). We find that osmotic stretch alone fails to activate the kinase activity of mTORC2 but can significantly amplify the chemoattractant-stimulated kinase activity of the complex. PI3K activity is necessary and sufficient to amplify the kinase activity of mTORC2 upon osmotic stretch (**Fig. 5**). Our results suggest that a combination of biochemical inputs and mechanical stretch can combine to trigger mTORC2 activity to inhibit actin assembly in protrusions (which contain the biochemical co-input PIP_3_). An attractive hypothesis is that the kinase activity of mTORC2 could have a different spatial range to independently regulate myosin-based contractions in regions not constrained by PIP_3_, though future imaging-based activity probes will be necessary to address the spatial logic of mTORC2 activation during cell polarity and movement.

## MATERIALS & METHODS

### Cell Lines and Culture

Cell culture was performed as described previously (Graziano et al., 2019). HL-60 cells were cultured in RPMI-1640 with 25 mM HEPES and L-glutamine (Corning) with 10% (v/v) heat- inactivated FBS (Gibco) and 1X penicillin/streptomycin (Gibco) and maintained at 0.2–1.0 million cells/ml. HL-60 cells were differentiated with 1.5% DMSO (Sigma-Aldrich) at a starting density of 0.2 million/ml in growth media for 5 days to obtain neutrophil-like dHL-60 cells. All experiments were performed with dHL60s unless otherwise stated. HEK-293T cells (to generate lentivirus for transducing HL-60s) were cultured in DMEM (Gibco) with 10% (v/v) heat-inactivated FBS. All cells were cultured in a 37°C/5% CO_2_ incubator and routinely monitored for mycoplasma contamination.

### Plasmids

Plasmids were constructed using standard molecular biology protocols. DNA segments were PCR-amplified and cloned into a pHR lentiviral backbone and driven by promoter from spleen focus-forming virus (SFFV) via standard Gibson assembly between the MluI and NotI restriction sites. A construct for Rac biosensor PakPBD-mCherry was generated from PakPBD-mCitrine (Graziano et al., 2019) by switching the DNA encoding the fluorescent tag. The RhoA biosensor uses the AnillinAHPH domain (Piekny and Glotzer, 2008) and was obtained from Addgene (plasmid # 68026). The EGFP-AnilinAHPH was sub-cloned into pHR lentiviral backbone using similar sites and strategy as mentioned above. We modified our previously reported Opto-PI3K module from red-light sensitive Phy/PIF (Graziano et al., 2017) to blue-light sensitive iLID (Guntas et al., 2015). The Opto-PI3K module consists of two constructs which need to be co-expressed for exogenous PIP_3_ generation - Construct A: iLiD-BFP-CAAX membrane anchor; Construct B: iSH2-EGFP-SspB scaffold for PI3-Kinase recruitment. Both constructs were assembled by Golden Gate-based cloning from a library of individual components from an extendable Mammalian Toolkit (Fonseca et al., 2019) into the pHR vector backbone with EF1α promoter (for Construct A) or SFFV promoter (for Construct B). Guide RNAs (gRNAs) with homology to exon 2 of RICTOR (5’ GTCCCGCTGGATCTGACCCG 3’) and exon 2 of MAPKAP1/mSin1 (5’ AGTCAGTCGATATTACCTCA 3’) were selected using the CRISPR design tool in Benchling (https://benchling.com/) and cloned into the LentiGuide-Puro vector (Addgene plasmid #52963) as previously described (Sanjana et al., 2014). The pHR vector used to express human codon–optimized *Streptococcus pyrogenes* Cas9-tagBFP was as previously described (Graziano et al., 2017).

### Lentiviral transduction of HL-60 cells

HEK-293Ts were plated in 6-well plates (Corning) and grown to 70–80% confluency. Cells were transfected with 1.5 µg of the pHR plasmid along with two plasmids containing the lentiviral packaging proteins (0.167 µg of pMD2.G and 1.3 µg of p8.91) with TransIT-293 (Mirus Bio). After 2–3 d of transfection, lentivirus-containing supernatant was collected, filtered with a 0.45-µm filter (EMD Millipore), and concentrated 40-fold with Lenti-X Concentrator (Takara). The concentrated lentivirus was used immediately or stored at −80°C. HL-60 cells were transduced by overnight incubation of 0.3 million cells with 4 µg/ml polybrene and ∼130 µl of concentrated virus. Cells expressing desired transgenes were isolated by culturing in growth media supplemented with puromycin (1 μg/mL) or using fluorescence-activated cell sorting (FACS) as appropriate (FACS Aria3, BD Biosciences).

### Generation of knockout cell lines using CRISPR/Cas9

RICTOR and MAPKAP1(mSin1) HL-60 KO cell lines were generated and validated as previously described (Graziano et al., 2019, 2017). Wildtype HL-60 cells were transduced with a puromycin- selectable vector containing the gRNA sequence targeting the gene of interest. Following puromycin selection, cells were transduced with a *S. pyrogenes* Cas9 sequence fused to BFP. Cells expressing high levels of Cas9-BFP were isolated with FACS, after which a heterogeneous population was obtained and assessed by sequencing of the genomic DNA flanking the Cas9 cut site. Cells were then diluted into 96-well plates at a density of approximately one cell per well in 50% (vol/vol) filtered conditioned media from a healthy culture, 40% (vol/vol) fresh HL-60 media, and 10% (vol/vol) additional heat-inactivated FBS. Clonal cell lines were expanded and validated by genomic DNA sequencing to infer the indel distribution and immunoblots to assay loss of protein expression. Clonal lines for Rictor KO and mSin1 KO were further assayed to check the residual mTORC2 kinase activity using phospho-Akt immunoblots.

### Cellular treatments and perturbations

Neutrophil-like dHL60 cells were activated with chemoattractant formyl-Met-Leu-Phe (fMLP; Sigma) at effective final concentration of either 25 nM (for imaging based assays) or 100 nM (for cellular biochemistry). Acute increase in membrane tension and stretching was done by adding equal volume of hypotonic buffer (H_2_O + 1 mM MgCl_2_ + 1.2 mM CaCl_2_) as described earlier (Graziano et al., 2019; Houk et al., 2012). mTOR kinase activity was inhibited by treating cells with 10 μM KU-0063794 (Selleckchem) for 30-45 mins in plain RPMI-1640. PI3Kinase activity was inhibited by incubating cells with 1 μM PIK-90 for 30-45 mins in plain RPMI as described earlier (Van Keymeulen et al., 2006). ROCK inhibition was done by treating cells with 20 μM Y-27632 for 20 min in plain RPMI prior to the imaging assay (Graziano et al., 2019).

### Cellular Biochemistry

#### mTORC2 activity (phospho-Akt) and Rac Activity (phospho-Pak) assay

These assays were performed as described earlier (Graziano et al., 2017). Cells (dHL-60; 5 days after differentiation) were serum starved to reduce basal signals by incubation in plain RPMI for 30-45 min at 37°C/5% CO_2_ at a density of ∼ 1.5 million cells/ml. All time courses were performed at room temperature of 24°C. For chemoattractant-based time courses (e.g., Fig 3B, 5B) cells were activated at final effective fMLP concentration of 100 nM, and samples were collected at indicated time points by mixing 0.5 ml of cells with 0.5 ml ice-cold stop solution (20% TCA, 40 mM NaF, and 20 mM β-glycerophosphate). Samples were incubated in TCA at 4°C for at least 1h, after which proteins were pelleted, washed once with 0.75 ml ice-cold 0.5% TCA, and solubilized in 2x Laemmli sample buffer (Bio-Rad Laboratories). For pAkt assays (S1 Fig C and Fig 5E) cells were assayed in absence (basal level) or 3 minutes after 100 nM fMLP addition, as pAkt signals peaked at 3 min in our assays. Optogenetic stimulation of Opto-PI3K dHL-60 cells (Fig 5H) was done with the following modifications to the protocol above. Serum starved cells were placed in wells of a clear-plastic 24 well plate (Corning) and placed about 1 cm above a blue (450 nm) LED array. A ND4 filter (Sioti) was inserted between the cells and the LED light source to attenuate the illumination to 1mW. Cells were illuminated for 3 min following which the LED was switched off and chilled TCA was added to prepare samples as described above. Samples were then analysed by immunoblots. Quantification of these assays were done by calculating the ratio of band intensities of phospho-Pak (or Akt) to the total-Pak (or Akt). These values were then normalised to the peak values obtained for wildtype control in the time-series (generally at 1 min for pPAK and 3 min for pAkt).

#### Immunoblot assay

Western blotting was done as previously described (Graziano et al., 2017). Briefly, protein content from at least 0.75-1 million HL60 cells was extracted by chilled TCA precipitation and resuspended in 2x Laemmli sample buffer. Protein samples were separated via SDS-PAGE, followed by transfer onto nitrocellulose membranes. Membranes were blocked for at least 1 hr in a 1:1 solution of TBS (20 mM Tris, 500 mM NaCl, pH 7.4) and Odyssey Blocking Buffer (LI-COR) followed by overnight incubation at 4 °C with primary antibodies diluted 1:1000 in a solution of 1:1 TBST (TBS + 0.2% w/v Tween 20) and Odyssey Blocking Buffer. Membranes were then washed 3x with TBST and incubated for 45 min at room temperature with secondary antibodies diluted 1:10,000 in 1:1 solution of Odyssey Blocking Buffer and TBST. Membranes were then washed 3x with TBST, 1x with TBS and imaged using an Odyssey Fc (LI-COR). Analysis was performed using Image Studio (LI-COR) and Excel. The following primary antibodies were used for the study; phospho-PAK1 (Ser199/204)/PAK2 (Ser192/197) (Cell Signaling #2605), PAK2 (3B5) (Cell Signaling #4825), phospho-Akt (Ser473; D9E) XP (Cell Signaling #4060), Akt (pan; 40D4; Cell Signaling #2920S), Rictor (Bethyl #A300-458A), mSin1 (Bethyl # A300-910A), and GAPDH Loading Control Antibody GA1R (ThermoFisher). Secondary antibodies IRDye 680RD Goat anti-Mouse (LI-COR) and IRDye 800CW Goat anti-Rabbit (LI- COR) were used.

### Transwell Assays

These assays were performed as previously described (Diz-Muñoz et al., 2016). Briefly 0.3 million dHL-60 cells were stained with 5ul/ml DiD (Life Technologies) and plated on the upper chamber of the 24-well format HTS FluoBlokTM Multiwell Insert System (3 μm pore size; BD Falcon) in RPMI without phenol red (Life Technologies) with 2% FBS. Cells were allowed to migrate towards the bottom well containing 20 nM fMLP for 1.5 hr at 37°C. The migrated cells were measured by fluorescence from the bottom of the insert using FlexStation 3 Microplate Reader (Molecular Devices). Migration index was calculated by dividing the amount of signal in the sample well by the signal in a well in which 0.3 million cells were plated in the bottom well. The dataset was normalised by the peak value obtained for wildtype cells (usually observed at 60 min).

### Imaging Assays and Data Analysis

#### Microscopy hardware

We used a spinning-disk confocal microscope for all imaging data presented here. The setup comprised of a Nikon Eclipse Ti inverted microscope with following objectives (Plan Apo 10x/0.45 NA, 20x/0.75NA, 60x/1.40 NA, 100x/1.4 NA; Nikon), Yokogawa CSU-X1 spinning-disk confocal, Prime 95B cMOS camera (Photometrics), 4-line laser launch (405, 488, 561 and 640 nm laser lines; Vortran) and environmental control (37°C/5% CO_2_; Okolab).

#### Preparation of dHL-60s for microscopy

Imaging based assays with dHL-60 cells were performed using 96-well #1.5 glass-bottom plates (Brooks Life Sciences). The wells were coated with a 100 µL solution of 10 µg/mL porcine fibronectin (prepared from whole blood) and 11 mg/mL bovine serum albumin (BSA, endotoxin- free, fatty acid free; A8806, Sigma) dissolved in Dulbecco’s Phosphate Buffered Saline (DPBS; 14190-144, Gibco) and incubated for 30 min at room temperature. Fibronectin solution was then aspirated, and each well was washed twice with DPBS. dHL-60 cells in growth media were pelleted at 300xg for 5 min, resuspended in 100 µL imaging media (RPMI1640 with 0.5% FBS and 1 nM fMLP), plated in each well and incubated (37°C/5% CO_2_) for 10 minutes for cells to adhere.

#### F-actin staining

Cells were prepared and plated in 96-well glass bottom plate as described above and stimulated with 50 nM fMLP. At desired timepoints (usually 1min and 5 min), an equal volume of 2x fixation buffer (280 mM KCl, 2 mM MgCl2, 4 mM EGTA, 40 mM HEPES, 0.4% bovine serum albumin (BSA), 640 mM sucrose, 7.4% formaldehyde (w/v), pH 7.5) was added to each well and incubated for 15 mins at room temperature (RT). The fixation buffer was then removed from each well and cells are washed once with intracellular buffer (140 mM KCl, 1 mM MgCl2, 2 mM EGTA, 20 mM HEPES, 0.2% BSA, pH 7.5). Following fixation, cells were treated with staining buffer (intracellular buffer + 5 ul/ml Alexa647-Phalloidin (Invitrogen) + 0.2% Triton X-100) for 30 mins in dark at room temperature. Cells were finally washed gently to remove excess staining buffer and 200 ul of intracellular buffer mixed with nuclear dye NucBlue (Thermofisher) was added to each well.

#### Immunofluorescence

Cells were prepared, plated, and fixed as above following which cells were incubated with permeabilization buffer (intracellular buffer and 0.2% Triton X-100) for 20 mins at RT. Cells were then blocked (3% BSA and 1% normal goat serum in permeabilization buffer) for at least 1h at RT. Cells were washed and then incubated with primary antibody diluted in blocking solution for at least 2h at RT or overnight at 4°C. Cells were washed and then incubated with secondary antibody for 45 min-1h at RT. Finally, cells were washed 2x with permeabilization buffer and 1x with intracellular buffer before adding fresh intracellular buffer mixed with NucBlue to each well for imaging. Phospho-Myosin Light Chain (Ser19) primary antibody (rabbit; Cell Signaling #3671) was used at 1:200 dilution with secondary antibody Goat anti-Rabbit IgG Alexa 488 (Invitrogen #A-11034) at 1:1000 dilution.

#### Image analysis for fixed preparations

All image analysis to measure the levels and distribution of F-actin and pMLC was performed with Fiji-ImageJ. Briefly, raw images comprising z-stacks of several fields of cells obtained from the Nikon spinning disk confocal (ND2 format) were imported into Fiji. Before quantification, the image were background and flat field corrected using the background and flat-field fluorescence values estimated individually for all the different emission channels. Using the z-project tool in Fiji the corrected z-stacks were converted into maximum intensity projection for visualisation and sum-intensity projections for quantification of fluorescence. Intensity thresholds were estimated from the pixel intensity histogram and uniformly applied to sum-intensity projections to identify cell bodies and ‘measure stack’ function was used to measure intensity value for the whole field. Several fields (∼ 20) with 10-15 cells each were pooled from multiple independent experiments to quantify the fluorescence levels of F-actin and pMLC reported here and data was represented as box-plot to show the entire distribution of measured values. To quantify the width of the F- actin rich zone at cellular fronts, a 5 μm line ROI (10 pixel wide; 2 μm) was drawn from the front of the cell to obtain the intensity v/s length profile for each cell. Several cells (∼20-30 for each condition) were measured and their respective profiles were averaged across the whole dataset. The averaged intensity profile was fitted to a Bi-gaussian distribution of skewed peak to account for the overall width of the F-actin rich zone. To visualize the 3D distribution of axial z-protrusions of cells, ROIs of individual cell z-stacks were imported into the 3D visualisation software UCSF- ChimeraX and intensity thresholded to highlight the F-actin structures and the cell nuclei. Fraction of cells with axial protrusions was calculated by visual inspection and counting of all cells with 3D-projections and the total number of cell nuclei in the entire field of F-actin stained cells.

#### Under-agarose preparation of HL-60s for imaging

Cells were prepared using standard under-agarose preparation techniques as described previously (Bell et al., 2018; Brunetti et al., 2022). Briefly, a solution of 2% w/v low melt agarose (GoldBio) dissolved in RPMI1640 was prepared by heating the solution gently in a microwave. The solution was placed in a water bath at 37°C to cool down before adding to cells. In the meantime, dHL-60 cells were spun down at 300 x g for 3 mins and resuspended in plating media (RPMI + 2% FBS) at a concentration of 1 million/ml. 5 μl of cells were placed in the centre of a circular well (96 well plate; Greiner Bio-one) and allowed to settle for 5 min at RT. Agarose solution (195 μl) was then slowly dispensed into the well directly on the top of the cells. This allowed even deposition of agarose which congeals over 10-20 min at RT, following which the wells were monitored quickly under a standard tissue culture brightfield microscope and moved to the microscope to equilibrate at 37C for another 20 mins prior to imaging.

#### Cell Motility Assays

Single cell motility assays were performed with cells plated in an under-agarose preparation (to mimic in-vivo like confined environment) or on fibronectin coated glass (standard unconfined 2D motility). For both types of motility assays, 0.1 million cells were labelled using 1μM CellTracker Green or Orange (Thermo Fisher) in plain RPMI for 10 min at 37C. Labelled cells were spun down from labelling mix and washed once with plain RPMI and resuspended in imaging media (RPMI1640 + 0.5% FBS + NucBlue nuclear marker) and were either plated on fibronectin coated glass (as described above) or processed for under-agarose preparation and plates were moved to the microscope preheated to 37C for imaging. Cells were stimulated with uniform fMLP concentration of 25nM. To allow tracking of several cell nuclei, imaging was done using a low magnification objective (10x or 20x) to capture a larger field of view and imaged once every 30sec for over 20-30 min.Tracking of cell nuclei was performed using the Fiji plug-in TrackMate (Tinevez et al., 2017). Cell nuclei tracks were filtered for desired property (duration of at least 10 min, high quality, not undergoing collisions) to obtain the coordinates of movement. Track features (like speed, persistence, displacement) were computed with inbuilt tools of Trackmate.

#### Biosensor imaging

To measure the extent of temporal coordination of front-back polarity, cells expressing both the Rac (PakPBD-mCherry) and RhoA (EGFP-Anilin-AHPH) biosensors were prepared and plated under agarose (2% w/v in RPMI1640) and stimulated with uniform chemoattractant (25 nM fMLP). Cells were imaged on spinning disk confocal at high spatial resolution (60X or 100X objective) with fast sequential acquisition (exposure time 100 ms) for 3 minutes at frame interval of 3 sec using a similar strategy as previously described (Tsai et al., 2019). The raw images were background and flat-field corrected, and full images were manually cropped to a ROI with a single cell.

#### Analysis for spatial distribution of polarity

Spatial profiles of biosensor intensity were obtained from a line ROI (10 pixel wide; 1.5 μm) starting at the leading edge and extending to the uropod to record the intensity v/s length traces for both Rac and RhoA biosensor. Pearson’s correlation coefficient between the two traces were computed for several such individual cells (at least 20-30, pooled from independent experiments) and plotted as box-plots. This metric provided an intuitive method to assess the front-back correlation (or lack of it in wildtype cells) of spatial distribution of polarity signals. To assess the lateral correlation, another line ROI was drawn near the centre of the same cell perpendicular to the front-back axis. Intensity profiles for both biosensors were recorded, and correlation coefficients were calculated.

#### Analysis for temporal coordination of polarity

Cells which touch or collide with a neighboring cell were ignored from this analysis as they presented challenges to segmentation and further analysis. Single cells were then segmented by smoothing and intensity-based thresholding for each of the two emission channels for the biosensor intensities for every frame of the image sequence. The resulting binary images were then combined by taking the union of the two segmentations. Further analysis was done for sequence of frames where the segmented edge of the cell does not touch the boundary of the cropped image ROI. Using the consensus binary image described above, the weighted centroid of biosensor fluorescence intensity was calculated for each channel across frames. The distances between the coordinates of these centroids were calculated at each frame and the coefficient of variation (CV) of this series was calculated for each cell. CV provides a normalized metric for fluctuations in the inter-centroid distance over the course of the timeseries and can be readily compared for cells of varying sizes or across different genetic background (say, wildtype or Rictor KO cells). The full analysis code is available on GitHub (https://github.com/orgs/weinerlab/repositories).

## ACKNOWLEDGEMENTS

We thank all members of the Weiner laboratory for stimulating discussions and Brian Graziano for experimental assistance; Ben Winer and Madhuja Samaddar for critical reading of the manuscript. We thank Kevan Shokat for providing PIK-90. This work was supported by American Heart Association Postdoctoral Fellowship AHA-18POST33990156 (S Saha), UCSF Program for BreAkthrough Biomedical Research Independent Postdoctoral Grant (S Saha), NSF Graduate Research Fellowship Program (Grant No. 1650113, JP Town), NIH Grant GM118167 (OD Weiner), National Science Foundation/Biotechnology and Biological Sciences Research Council grant 2019598 (OD Weiner), the National Science Foundation Center for Cellular Construction (DBI-1548297).

The authors declare no competing financial interests.

## Author contributions

Conceptualization: S Saha, OD Weiner; Investigation and data curation: S Saha, JP Town; Formal Analysis: S Saha, JP Town; Reagents and Methodology: S Saha, JP Town; Funding acquisition : S Saha, JP Town and OD Weiner; Software: JP Town; Visualization and Figures: S Saha, JP Town; Writing and revising: S Saha (original draft) with edits and inputs from OD Weiner and JP Town.

## Supplementary Figures

**Fig S1.**
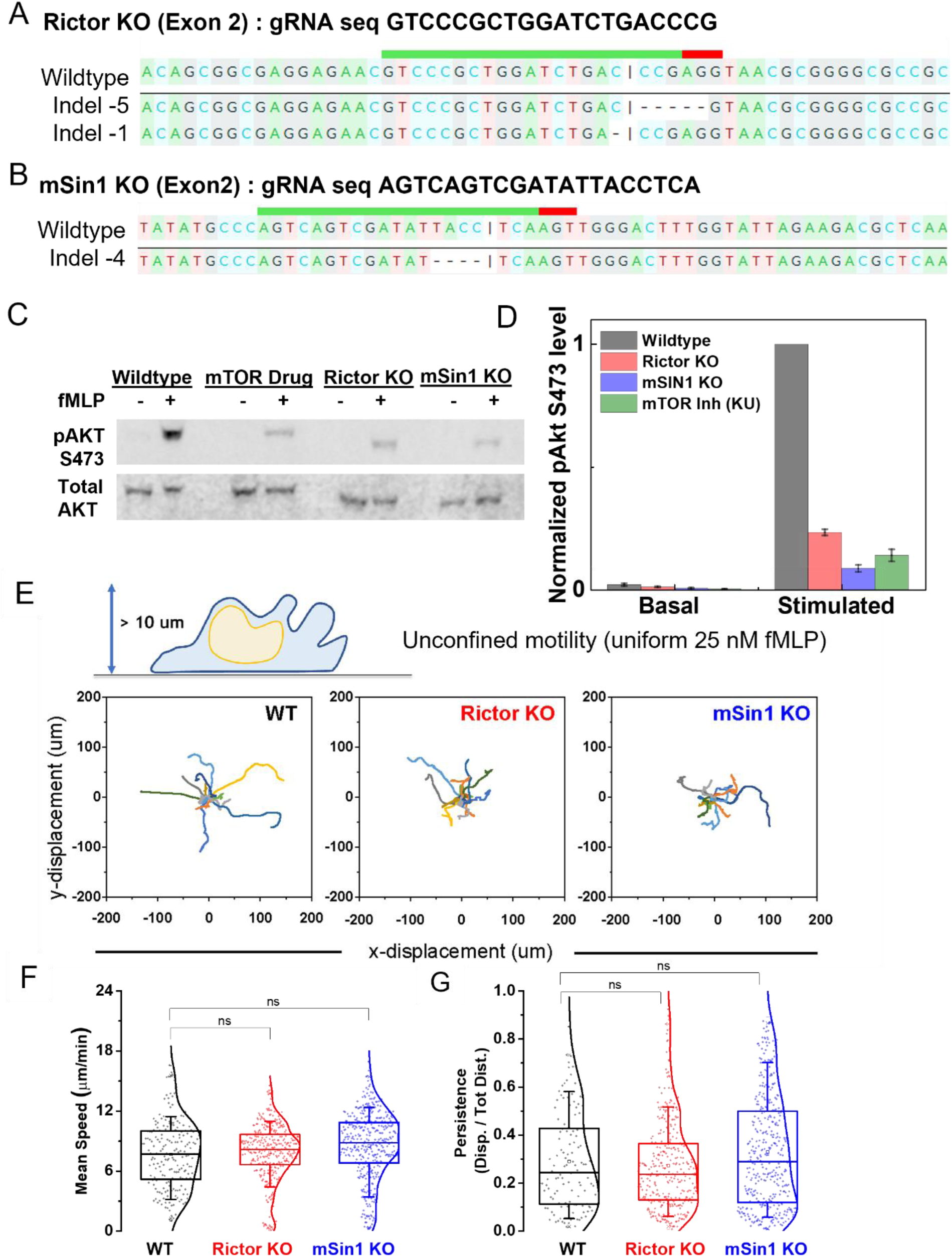
Gene-editing of knockout lines and functional validation of mTORC2 kinase activity and cell behavior in unconfined 2D motility assays. (A,B) Sequence validation to infer CRISPR indel edits in the clonal Rictor KO (A) and mSin1 KO (B) lines assayed here. The green bar above both sequences shows the gRNA target sequence. Both lines have deletions that lead to a frame shift, nonsense, and termination following Exon 2. (C, D) mTORC2 kinase activity assayed by immunoblots of phospho Akt and total Akt levels before (basal,-) and 3 min after chemoattractant (fMLP, +) addition. Representative western blots and quantification (D) shows significant loss of mTORC2 kinase activity for Rictor KO, mSin1KO and mTOR drug KU (assayed by pAkt immunoblots). Plots (D) show pAkt/totalAkt ratio (mean ± SEM from three independent trials) normalised with values obtained for the wildtype for each trial. (E) Schematic shows a neutrophil-like dHL60 cell undergoing unconfined motility on glass coated with fibronectin in presence of uniform chemoattractant fMLP. Randomly chosen representative tracks (15 each) of wildtype (WT), Rictor KO, or mSin1KO cells over a 12 min observation window; axes show x-y displacement in μm. (F, G) Box plots (with kernel smooth distribution curve) show mean speed (D) and persistence (E; ratio of displacement/distance) averaged over individual tracks. Both Rictor KO and mSin1 KO cells shows normal persistence and speed in unconfined 2D migration (p < 0.01; one-way ANOVA with Tukey-means comparison). N = 203 (WT), 338 (RictorKO) and 392 (mSin1KO) tracks from individual cells pooled across 2 independent experiments. For box plots, median is indicated by the line, inter-quartile range sets the box width and error bars indicate 10-90^th^ percentile.

**Fig S2.**
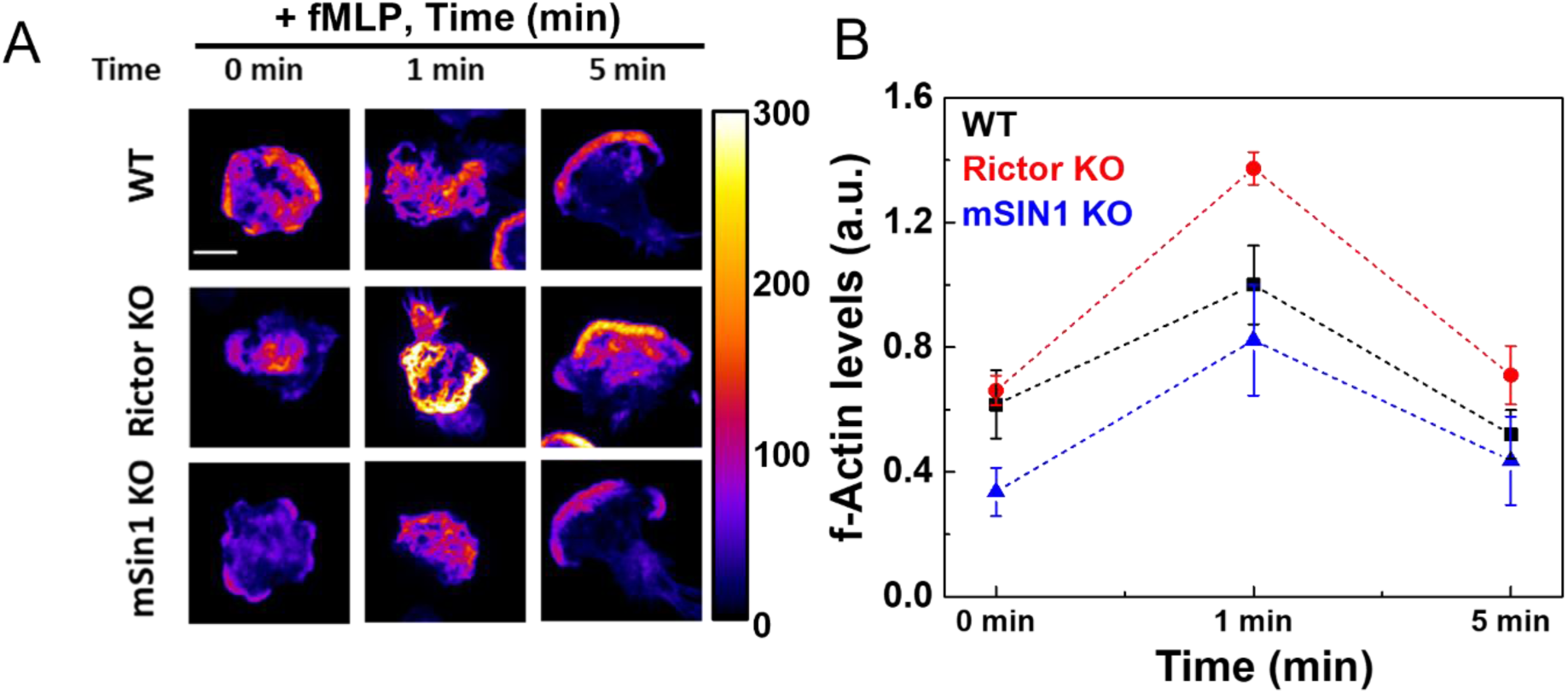
F-actin distribution and levels in wildtype, Rictor KO, and mSin1KO dHL60 cells upon chemoattractant stimulation. (A) Maximum-intensity projections of Alexa647-phalloidin stained F-actin obtained from z-stacks (10 μm deep) acquired on confocal spinning disk microscope for wildtype (WT), Rictor KO, or mSin1KO dHL60 cells, before and 1min or 5min after stimulation with 25nM fMLP (5 min images are duplicated from Fig 2B). (B) Normalised total F-actin levels (Mean ± SEM) from these experiments were quantified from confocal z-stacks (∼ 20 fields, at least 200-250 cells; pooled from 3 independent experiments) across each condition for the three genotypes. Mean F-actin intensity value at 1 min was used to normalise all conditions for each independent trial; scale bar is 10 μm.

**Fig S3.**
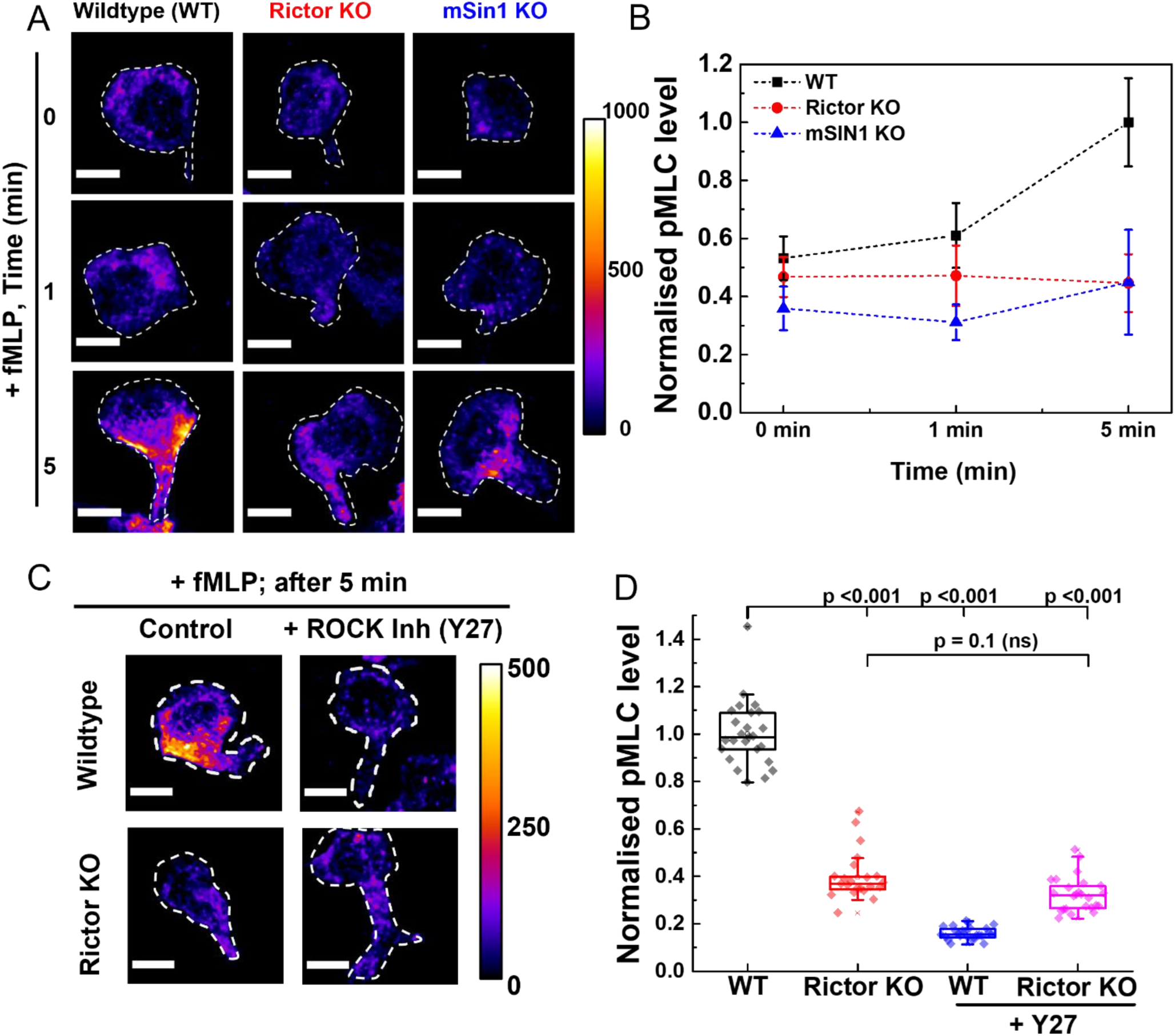
Kinase activity of mTORC2 stimulates myosin contractility. (A) Maximum intensity projections of pMLC immunostaining obtained from confocal z-stacks (9 μm deep) of wildtype (WT), Rictor KO, or mSin1KO dHL60 cells, before and 1 min or 5 min after stimulation with 25 nM fMLP (5 min images are duplicated from Fig 3D). All images are equally intensity scaled indicated Fire LUT; scale bar is 10 μm. (B) Normalised pMLC levels (Mean ± SEM) from these experiments were quantified from confocal z-stacks (∼ 15 fields, at least 150 cells; pooled from 2 independent experiments) across each condition for the three cell types. Mean pMLC intensity value at 5 min after stimulation was used to normalise all conditions for each independent trial. (C) pMLC immunostaining of wildtype or RictorKO cells either untreated or upon addition of 20 μM ROCK inhibitor Y27632 and stimulated with 25nM fMLP for 5 mins. Images show maximum intensity projections of z-stacks obtained from confocal z-stacks (9 μm deep) of cells; scale bar is 10 μm. (D) Box-plots of normalised pMLC level quantified from confocal z-stacks (∼20 fields, at least 250 cells; pooled from 3 independent experiments) across each condition. Y27 treatment leads to significant loss of pMLC levels in Wildtype cells comparable to levels seen in Rictor KO cells. Statistical significance was estimated at p<0.001 by one-way ANOVA with Tukey’s mean comparison test. Mean pMLC intensity value for WT (untreated) was used to normalise all conditions for each independent trial. For box plots, median is indicated by the line, inter-quartile range sets the box width, and error bars indicate 10-90^th^ percentile.

**Fig S4.**
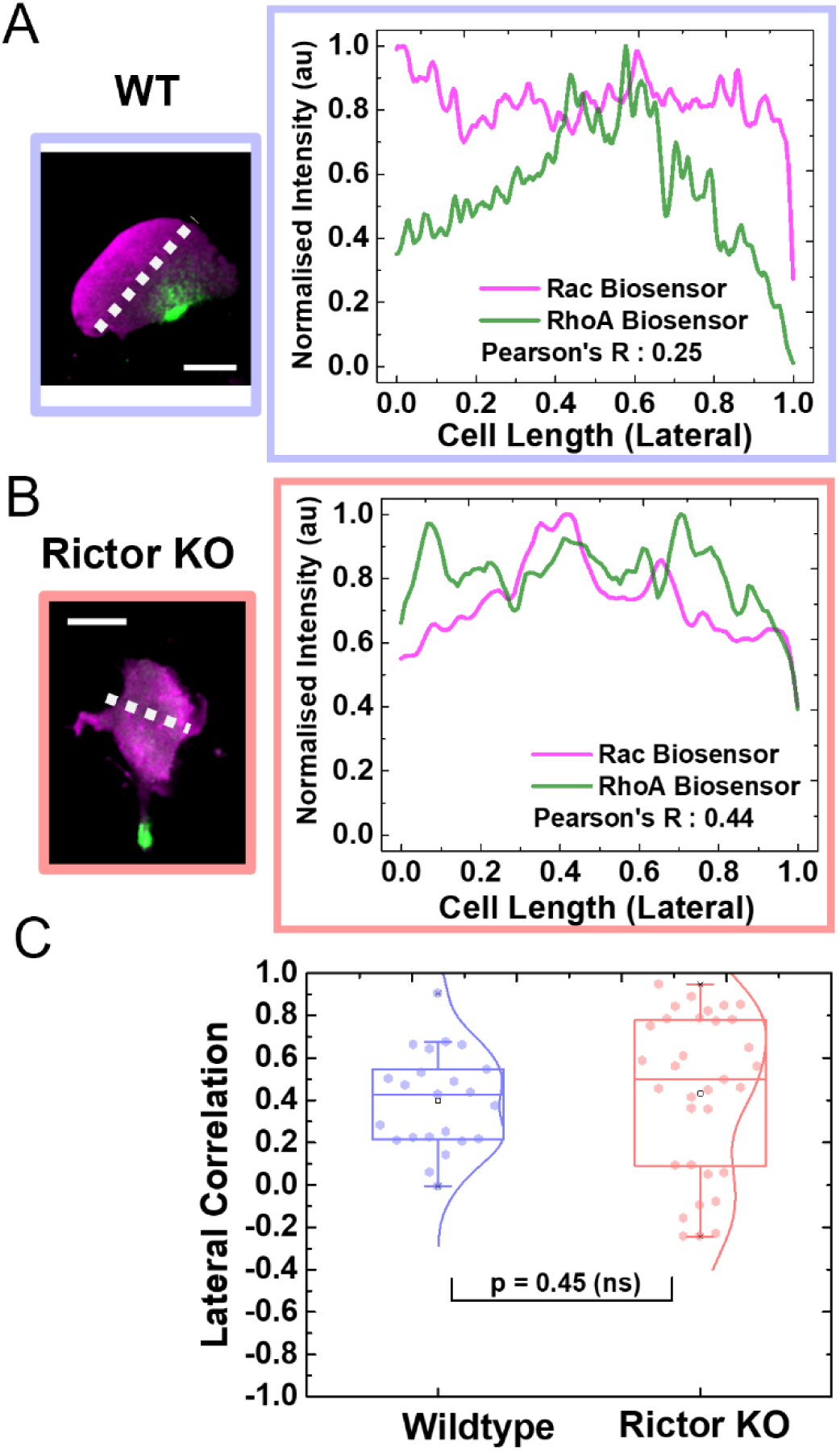
Polarity defects upon loss of Rictor/mTORC2 do not affect overall biosensor distrubution. A measure of correlation between the intensity profile of polarity signals across an axis orthogonal (or lateral) to the front-back axis provides an internal control of whether the overall biosensor distribution is skewed between wildtype and Rictor KO cells. We expect the front/back signals to exclude each other across front-back axis (as in Fig. 4) but not along the lateral axis. (A, B) Lateral intensity profile of same representative wildtype cell (as Fig 4A, WT panel) and Rictor KO cell (as Fig 4A,Rictor KO panel) expressing Rho* (green) and Rac* (magenta), with computed Pearson’s Correlation Coefficient (R) for each (R= 0.25 for WT; 0.44 for RictorKO); scale bar 10 μm. (C) Box-plots of lateral correlation for wildtype (WT, n=23 cells) and Rictor KO cells (n= 33 cells) pooled from 4 independent experiments. Wildtype and Rictor KO cells show similar distribution of correlation coefficient (p = 0.45; Mann-Whitney’s test), suggesting that overall distribution of the biosensors is not impaired in the lateral axis. For box plots, median is indicated by the line, inter-quartile range sets the box width, and error bars indicate 10-90^th^ percentile.

## Supplementary Videos

**Video 1.** Movie shows the ChimeraX 3D-reconstruction of representative example of Wildtype (left), Rictor KO (centre) and mSin1 KO (right) cell in the tilted *xz-plane* orientation to highlight the axial features of f-actin distribution; yellow-gold and blue represent F-actin and nucleus respectively.

**Video 2.** Movie of Wildtype (WT, Top) and Rictor KO (botton) cells co-expressing Rac biosensor (Pak- PBD-mCherry) and RhoA biosensor (EGFP-Anillin-AHPH) migrating under 2% agarose overlay. Movie shows the distribution of front (Rac*, magenta) and back (Rho*, green) polarity signals (Image, left) and the corresponding front (magenta arrow) and back (green arrow) polarity vectors. The cell centroid for each frame is indicated by the open circle and it’s displacement from the grid indicates overall cell movement.

## Notes

### Competing Interest Statement

The authors have declared no competing interest.

